# Supervised learning of protein melting temperatures: cross-species vs species-specific prediction

**DOI:** 10.1101/2024.10.12.617972

**Authors:** Sebastián García López, Jesper Salomon, Wouter Boomsma

## Abstract

Protein melting temperatures are important proxies for stability, and frequently probed in protein engineering campaigns, for instance for enzyme discovery and protein optimization. With the emergence of large datasets of melting temperatures for diverse natural proteins, it has become possible to train models to predict this quantity, and the literature has reported impressive performance values in terms of Spearman rho. The high correlation scores suggest that it should be possible to accurately predict melting temperature changes in engineered variants, and to reliably identify naturally thermostable proteins. However, in practice, results in these settings are often disappointing. In this paper, we explore this apparent discrepancy. We show that Spearman rho over cross-species data gives an overly optimistic impression of prediction performance, and that this metric reflects the ability to distinguish global differences in amino acid composition between species, rather than the specific effects of genetic variation. We proceed by investigating whether cross-species training on melting temperature is beneficial at all, compared to training specific models for each species. We address this question using four different transfer-learning approaches and a fine-tuning procedure. Surprisingly, we consistently find no benefit of cross-species training. We conclude that 1) current models for supervised prediction of melting temperature perform substantially worse than the literature suggests, and 2) that reliable transfer across species is still a challenging problem. An implementation of this work is available at https://github.com/deltadedirac/thermocontrast_tm

## Introduction

Reliable prediction of protein thermal stability is a long-standing challenge in protein engineering, particularly for enzyme optimization, where it is a strong indicator for functional integrity under thermal stress. Special focus has been on the prediction of the *changes* in stability induced by mutations, also referred to as ΔΔ*G*. Over the last decades, a long list of algorithms have been developed for this purpose, ranging from early models such as FoldX Delgado et al. [2019] and Rosetta Park et al. [2016], Frenz et al. [2020], Sora et al. [2023], to deep learning approaches based on convolutional and graph neural networks (Boomsma and Frellsen [2017], Li et al. [2020], Blaabjerg et al. [2023]), and most recently approaches building on large pre-trained models Dieckhaus et al. [2023].

The task of predicting *absolute* stabilities is generally considered more difficult, although some recent success in this area has been reported using (unsupervised) inverse-folding models Cagiada et al. [2024]. Supervised learning in this area has traditionally been challenging due to lack of large scale data, but new high-throughput experimental studies are beginning to provide sufficiently large, consistent datasets to make this feasible. An example is melting temperature data, which is a useful proxy for absolute stability, with the advantage that it can be measured reproducibly in high-throughput assays and, unlike other measures of protein stability (e.g. assays for stability under specific conditions), is directly comparable across proteins. This makes it a valuable general optimization target, frequently correlating with stability to complex stress factors Sanchez-Ruiz [2010]. A recent large-scale dataset of melting temperature of 48,000 proteins across 13 species Jarzab et al. [2020], combined with standardized splits Dallago et al. [2021], has made the melting temperature prediction problem accessible to the machine learning community. This has led to a flurry of new prediction methods, typically using pre-trained foundation models as the basis for supervised training. Two common approaches are 1) to train a *new* task-specific model using static embedding vectors from a pre-trained model as input, or 2) to *fine-tune* the foundation model on task-specific data, often using some parameter efficient technique to avoid overfitting. Several recent studies have reported impressive prediction performance, with Spearman correlation coefficients above 0.7 Dallago et al. [2021], Su et al. [2023], Sułek et al. [2024], Rodella et al. [2024], Pudžiuvelytė et al. (2024), Li et al. [2023], Yang et al. [2022].

Although progress in this area is highly welcome, the high performance values reported in the literature are somewhat at odds with our expectation about the difficulty of this prediction task. As mentioned, we would generally expect absolute stability prediction to be more challenging than relative stability prediction, where we do not see such high correlation coefficients. But more specifically, it conflicts with our own prior experience with melting temperature prediction in the protein engineering setting, where we have typically seen substantially lower performances than those reported. In this paper, we investigate the origin of this apparent discrepancy as a stepping stone towards a better understanding of the status of supervised learning performance in this domain.

The paper is organized as follows: We start by identifying the basis for the performance discrepancy described above – and show that much of the reported Spearman correlation arises as a consequence of the difference in melting temperature between species, rather than an ability to predict melting temperatures for individual variants. We thus confirm that species-specific melting temperature remains a challenging problem, with current methods displaying RMS errors of about 6 degrees, which are substantial, given the inner-species variance. Motivated by the large discrepancy between species, we then investigate whether there is any meaningful transfer of information between species, by comparing to models trained on a single species at a time. We conduct this analysis using a collection of different training approaches: 1) A simple transfer learning approach based on an ESM2 Lin et al. [2022] embedding, 2) a transfer-learning approach combining embeddings from sequence (ESM2) and structure-based (PiFold Hsu et al. [2022]) foundation models, 3) a transfer-learning procedure built on contrastive representation learning Zha et al. [2024], 4) a transfer-learning approach that explicitly anchors the results to a proxy for the optimal growth temperature of the species, and 5) a recently proposed standardized fine-tuning approach of the ESM2 model Schmirler et al. (2024). Our results demonstrate that across all these training procedures, we consistently see species-specific models outperforming globally trained models, despite the substantially smaller datasets. We conclude by discussing the implications of these results in terms of best practices for supervised melting temperature prediction.

## Results

We start by establishing a simple model architecture as a baseline for our subsequent analyses (Figure 1a). The model takes ESM2 embeddings Lin et al. [2022] as input, and produces melting temperature values as output. For simplicity, we use a classic transfer-learning setup with frozen embeddings (no fine-tuning). To aggregate the per-position embeddings into a single output we use a light attention layer, as this is known to outperform simple averaging over the length of the protein Stärk et al. [2021], Detlefsen et al. (2022). We analyze our performance on the Meltome Atlas dataset Jarzab et al. [2020], using the standard splits as provided in the FLIP benchmark Dallago et al. [2021]. Figure 2a (three left-most bars) shows that this baseline marginally outperform the results reported in the original FLIP paper, while providing somewhat lower performance than a recently published dedicated melting temperature predictor DeepSTABp Jung et al. (2023), demonstrating that our baseline implementation serves as a reasonable representative of the current capabilities in the field. See *Materials and Methods* for details.

**Figure 1:**
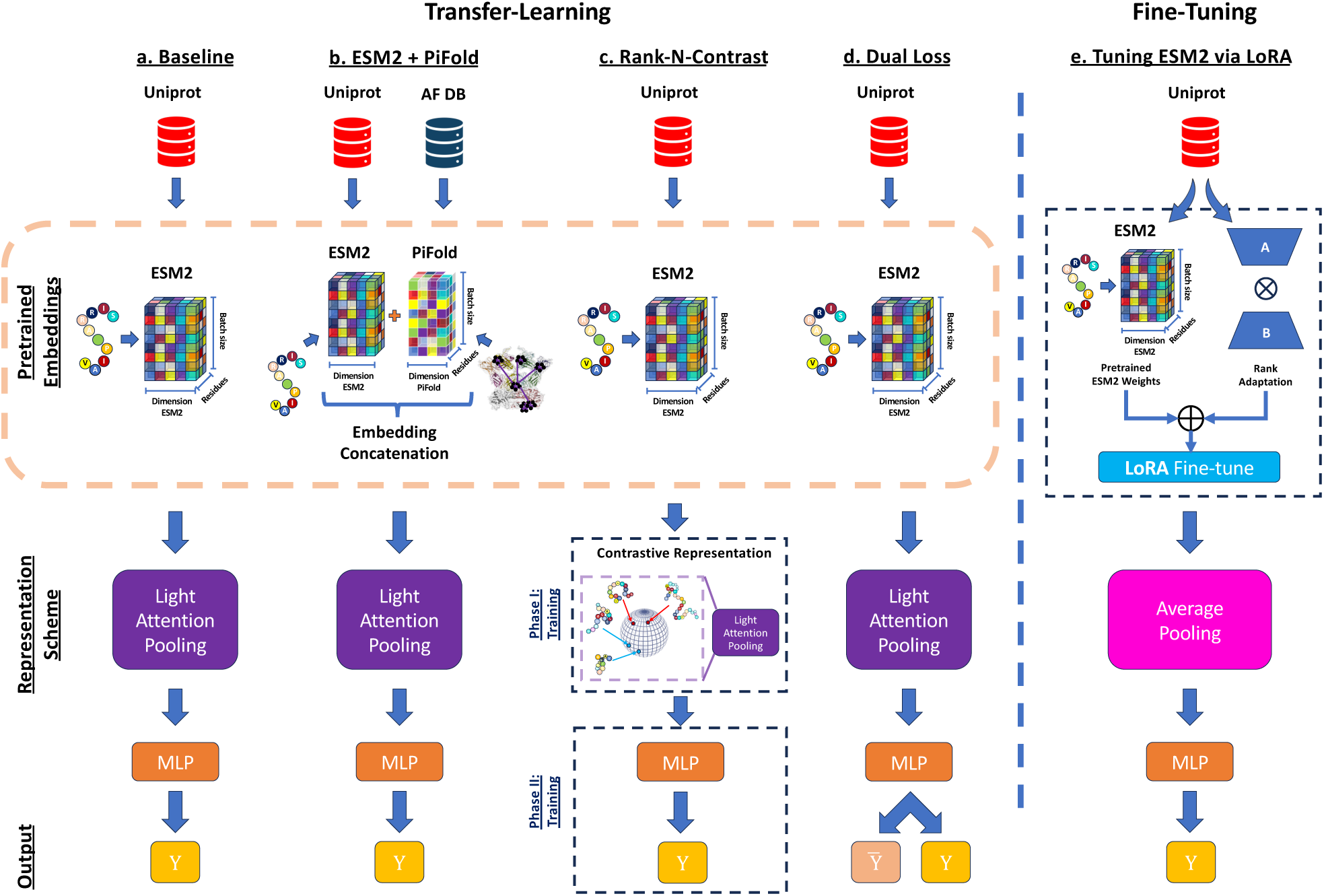
Five different prediction approaches, building on pretrained models. a) a simple baseline, using ESM2 Lin et al. [2022] embeddings pooled using light-attention; b) Similar to the baseline, but using a richer featurization consisting of a concatenation of ESM2 and PiFold Hsu et al. [2022]), an inverse-folding model; c) an contrastive representation-learning approach using the Rank-N-Contrast procedure; d) A dual-loss approach, where melting temperatures are predicted as offsets from the species mean; e) a LORA-base fine-tuning approach.

**Figure 2:**
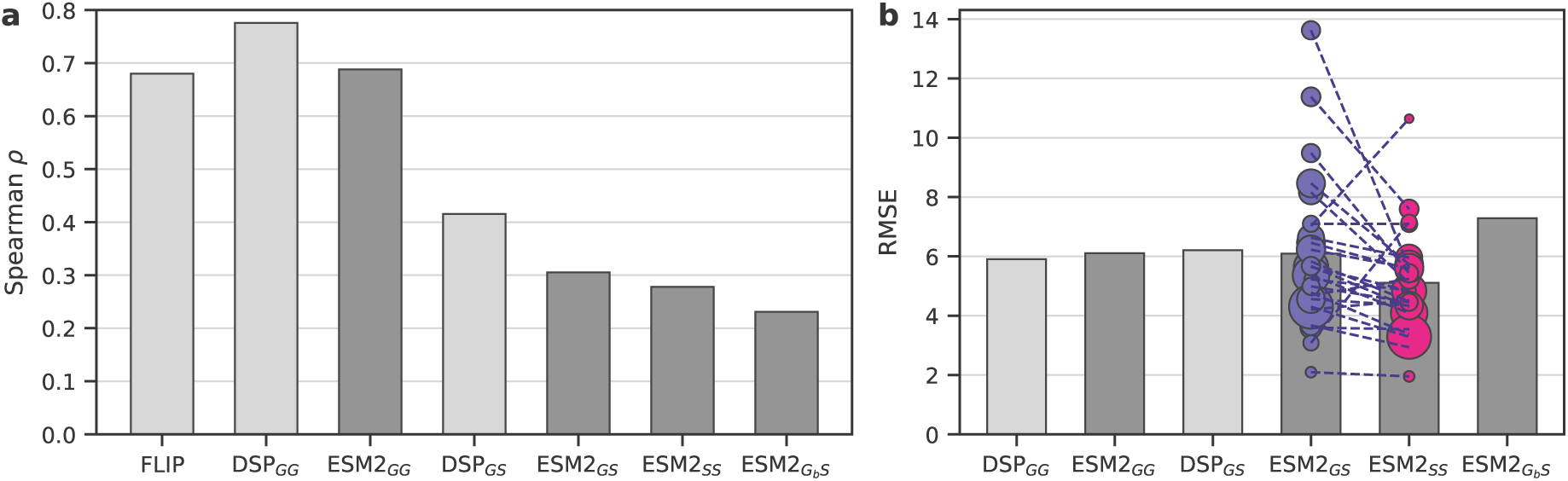
Performance of sequence-based embeddings to predict melting temperature across species. a) Performance measured in terms of Spearman rank correlation, b) Performance measured in terms of RMSE, showing the individual performances per species as circles, scaled by the size of the dataset. The G and S subscripts denote cross-species (*global*), and *specifies-specific*, respectively, with the first letter denoting the training scenario and the second denoting the testing scenario. For instance, *GS* denotes a model trained on all data, while being evaluated on each species. *G_B_* denotes a model that where the dataset was balanced by species during training (see *Methods*). Note how much of the correlation in the first two bars of the left plot arise simply from the fact that the correlation is measured across all species.

### Discrepancy explained: Spearman rho is a poor metric

The ESM2*_GS_* bar in Figure 2a is equivalent to the third (ESM2*_GG_*), but evaluates the Spearman correlation for each specifies *individually* and reports the average. The dramatic difference of this result compared to the three first bars explains the discrepancy discussed above. Clearly, much of the observed correlation found in the first two bars is due simply to global melting temperature differences between species, rather than the ability to determine melting temperature differences for different proteins within a species. This affect is apparent if we consider a scatter plot of the predictions coloured by species (Figure 3). This suggests that basing conclusions solely on the correlation of predictions with the ground truth may lead to misleading conclusions, and highlights the need to include additional metrics in the assessment of model performance. To illustrate, if we instead measure our performance in terms of root mean square error (RMSE), we find no notable discrepancy between the cross-species and per-species assessment of our model (Figure 2b).

**Figure 3:**
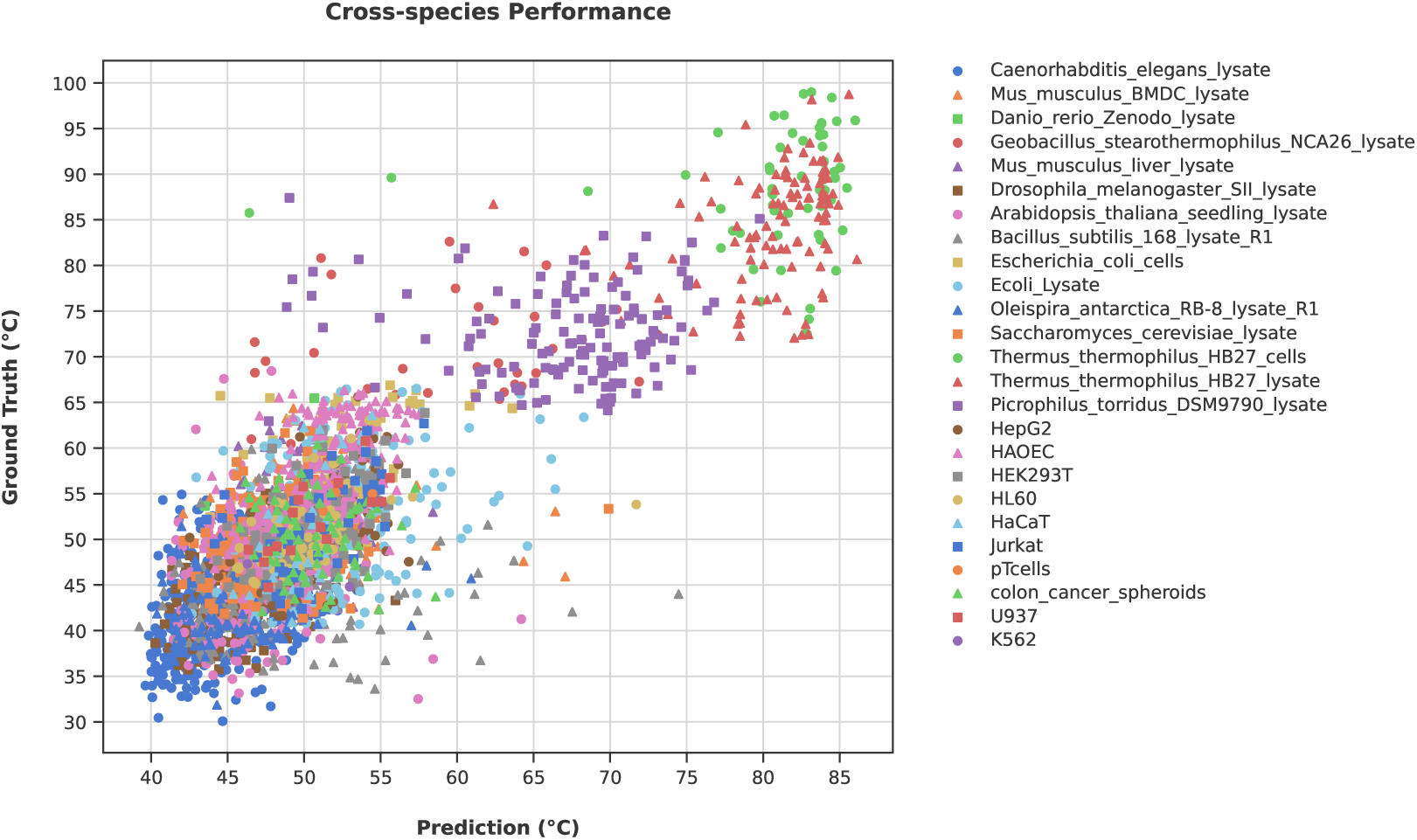
Scatterplot depicting the performance of the baseline cross-species model. Each point in the scatterplot is encoded by a unique marker and color combination, denoting the species of origin for the corresponding protein. This visualization complements the observations reported in Figure 2, highlighting that high Spearman rho arises as a consequence of cross-species variation, and that such analyses provide an overly optimistic picture of the prediction performance within species.

### Cross-species vs species-specific models

We have established that evaluating performance per-species aligns more closely with the objective of interest, namely assessing the impact of variation within species. A natural question is whether there is any cross-species transfer of information at all, or whether we would obtain better models if we simply trained separate models for each species. The training of cross-species vs species-specific models represents a trade-off: On one hand one might expect that cross-species models could benefit from larger datasets and could provide greater generalization capabilities; on the other hand, a specialized species-specific model might obtain a better fit with fewer parameters.

Assessing this question empirically with our baseline model on the FLIP dataset, we see that training species-specific models (ESM2*_SS_*, 2a, fourth column) seem to provide improved performance compared the globally trained model, suggesting that training larger cross-species models is not beneficial in this setting. We see this both in terms of species-level Spearman rho, RMSE, and as a general trend across the different species (Figure 2b, circles). We note that the lack of performance of the global model is not related to the imbalance of data set sizes between species. In fact, rebalancing the databaset during training leads to worse results (ESM2*_Gb S_*). Instead, the problem is likely related to the same behavior that led to the exaggerated performance of the global Spearman correlation: a Simpson-paradox like issue, where the model focuses on the global trends and disregard the local intra-species effects (Figure 3).

In order to establish whether this difference between cross-species models and species-specific models is a general phenomenon, we probe the effect on a range of different modeling and supervised training procedures. We will introduce them each in turn:

#### Approach 0: The baseline

As introduced above, this is a simple transfer learning approach from frozen ESM2 Lin et al. [2022] embeddings (Fig 1a).

#### Approach 1: Richer embeddings

An obvious candidate for improved performance is the source of the embedding itself. We do not expect substantial differences between the embeddings from different protein language models Michael et al. [2024], so we focus here on incorporating structural information by combining the ESM2 sequence embedding with embeddings obtained from the PiFold Hsu et al. [2022]) inverse folding model (Fig 1b).

#### Approach 2: Contrastive representation learning

Another appealing approach is *contrastive* learning, where a model is trained to enforce pairwise relationships between samples. We follow a recent approach, rank-N-contrast Zha et al. [2024], which uses a contrastive procedure to learn task-specific representations, which are subsequently used for the downstream regression phase (1c). The representation learning phase contrasts samples based on their ranking within the target space of continuous labels. For example, if two samples in a batch during training have the highest similarity in terms of melting temperature, they will be enforced to have the shortest distance in representation space. Since the original publication reported a beneficial role of data augmentation, we add a simple data augmentation step in the form of a small Gaussian perturbation of the embedding input vector (see *Materials and Methods* for details).

#### Approach 3: Anchoring to optimal growth temperature

Different organisms have different optimal growth temperatures (OGT), which reflects the ability of proteins and biomolecules to maintain their essential functions and structural integrity in different temperature environments. In an attempt to explicitly model this effect, we introduce a procedure that offsets the predictions relative to the optimal growth temperature (OGT) for the respective species. To simplify matters, we will assume that the optimal growth temperature has a constant offset to the average melting temperature of the proteins for that species, and therefore use the average *T_m_* of each species as a proxy for the OGT. The model now has two outputs, predicting the average for the species (shared for all proteins of the species), and a protein specific offset from this offset. The corresponding loss function is:

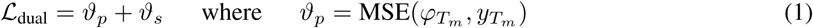

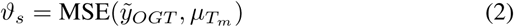

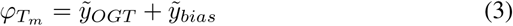

Here, *φ_Tm_* denotes the predicted *T_m_*, composed of two components: 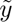*_OGT_* and 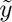*_bias_*, representing the two outputs from the final layer of the MLP *T_m_* predictor (see Figure 1 and section). The term 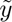*_OGT_* estimates the species average for each sample, while 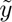*_bias_* indicates the deviation in degrees from this average. Additionally, *y_Tm_* denotes the ground truth melting temperature, and *µ_Tm_* is the average *T_m_* per species, serving as the reference OGT per species.

#### Approach 4: Parameter efficient Fine-tuning

As our final approach, we use a recently proposed protocol for parameter efficient fine-tuning of protein language models Schmirler et al. (2024). Here, rather than training a downstream model from scratch, the pre-trained model is instead adapted to improve performance on the task, by adding low-rank update matrices to the dense layers of the language model Hu et al. [2022]. Note that we for this protocol use simple average pooling instead of light attention pooling, in line with the published protocol Schmirler et al. (2024).

The results are presented in Figure 4, highlighting the differences across species and across methodological approaches. Overall, the different approaches show remarkably similar behavior, the fine-tuning approach perhaps showing slightly slower variance on the small datasets, while the rank-N-contrast is perhaps slightly less robust. The main take-away, however, is that the species-specific models seem to consistently outperform the cross-species models. We quantify these differences in Table 1, including paired t-test results that confirm significance of this conclusion in all 5 cases.

**Figure 4:**
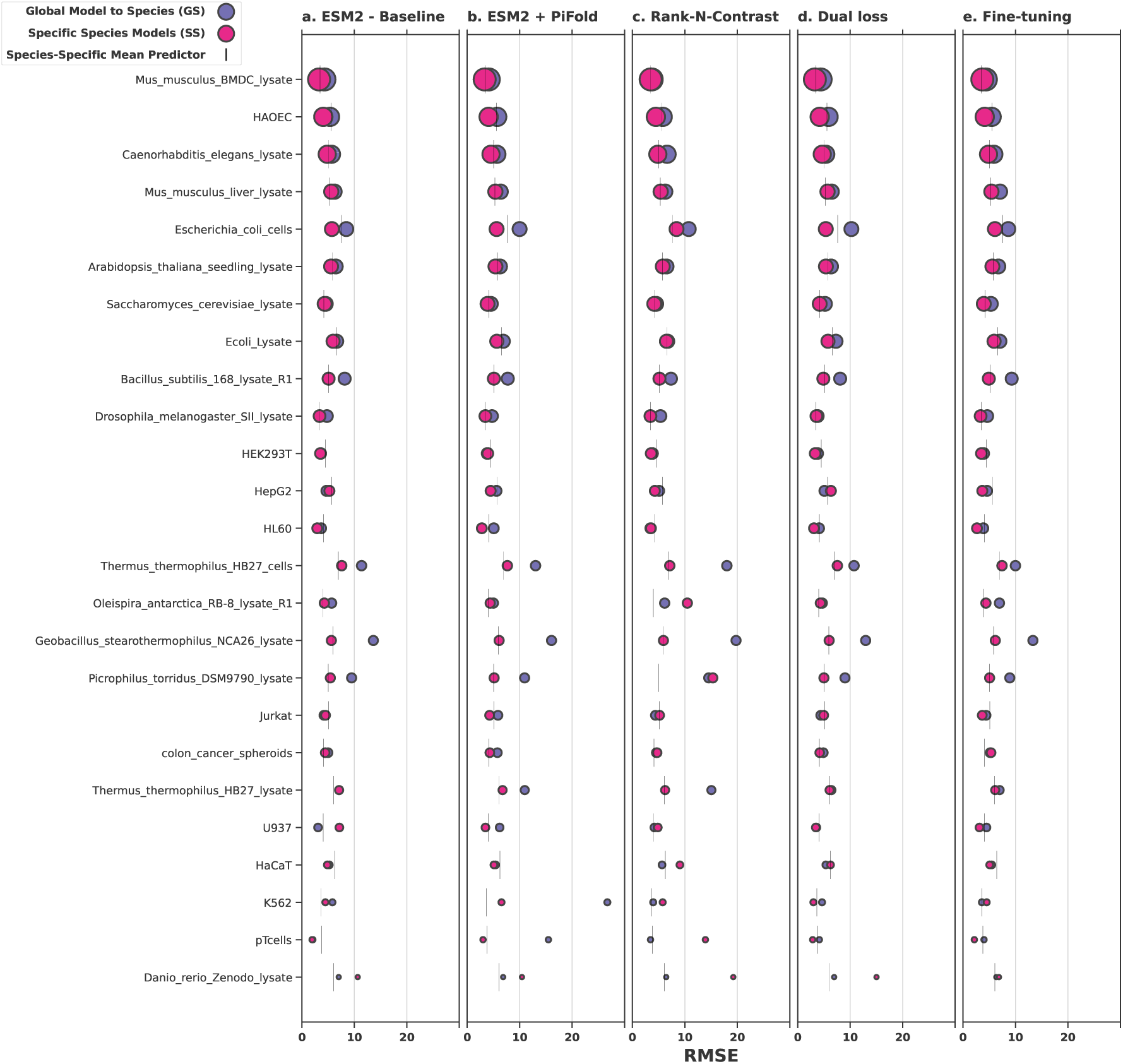
Performance of five methods across species, highlighting the difference between training a single model across species versus training separate models for each species. The size of the circles reflect the size of the datasets for each species. Results from a simple mean-prediction reference are included for reference.

**Table 1:** Overview of results in terms of an average root-mean-square-error (RMSE) and mean-relative-absolute- error (MRAE) for the individual datasets, for models trained on all species at once (cross-species) or for each species individually (species-specific). The compared methods include several transfer learning procedures with static embeddings, and a fine-tuning approach (see main text for details). A simple mean-averaging procedure is included as a reference. Row-wise paired t-tests were performed to assess whether the cross-species and species-specific methods were significantly different.

|  | Cross-species |  | Species-specific |  | t-statistic | p-value |
| --- | --- | --- | --- | --- | --- | --- |
|  | RMSE | MRAE | RMSE | MRAE |  |  |
| **Baseline** | $6.17 \pm 0.11$ | 8.9% (0.1pp) | $4.86 \pm 0.09$ | 7.2% (0.1pp) | 11.82 | 1.34e-31 |
| **ESM2+PiFold** | $6.77 \pm 0.12$ | 9.3% (0.2pp) | $4.71 \pm 0.08$ | 6.9% (0.1pp) | 16.21 | 8.97e-57 |
| **Rank-N-contrast** | $7.62 \pm 0.13$ | 10.4% (0.2pp) | $6.27 \pm 0.11$ | 8.4% (0.1pp) | 11.58 | 2.21e-30 |
| **Dual loss** | $6.20 \pm 0.11$ | 8.7% (0.1pp) | $4.81 \pm 0.09$ | 7.0% (0.1pp) | 12.68 | 5.83e-36 |
| **Fine-tuning** | $6.37 \pm 0.11$ | 9.2% (0.2pp) | $4.72 \pm 0.08$ | 6.9% (0.1pp) | 15.41 | 1.05e-51 |
| **Mean predictor** | $11.66 \pm 0.15$ | 15.2% (0.2pp) | $5.16 \pm 0.06$ | 8.0% (0.1pp) | 28.06 | 1.77e-154 |

**Figure 5:**
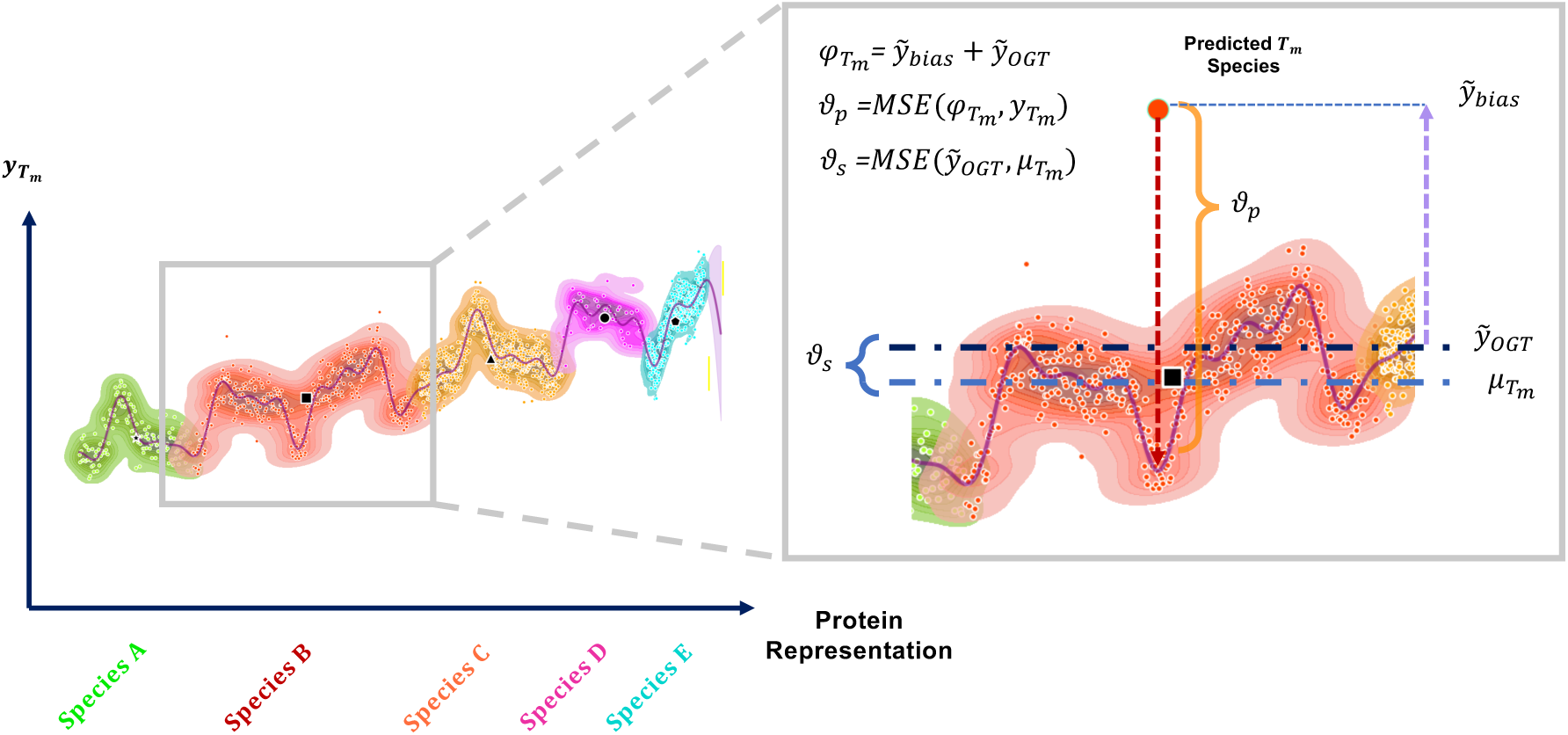
Schematic of the mechanism of action of the dual loss function, for simplicity depicting proteins along a 1-dimensional axis. In the graph, the densities represent high probability regions for the fit to *T_m_* for each protein within the training set. The density is color-coded to denote the species to which the points within that density belongs. The composition of the dual loss is depicted in the highlighted region. Here, *φ_Tm_* represents the predicted *T_m_*, calculated as the sum of 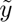*_OGT_* (predicted OGT) and 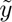*_bias_* (predicted bias), and *µ_Tm_* denotes the species-specific mean *T_m_*. The term *ϑ_p_* denotes the MSE between *φ_Tm_* and the ground truth *T_m_* (*y_Tm_*), while *ϑ_s_* represents the MSE between 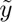*_OGT_* and the reference OGT (*µ_Tm_*). The dual loss is defined as the sum of *ϑ_p_* and *ϑ_s_*.

To give an impression of how good these predictions are in absolute terms, we also include an even simpler reference baseline for the species-specific models, where predictions merely consist of the mean melting temperature observed for that species in the training set (Figure 4a-e, vertical bars). It is remarkable that for many of the datasets, the gains obtained with state-of-the-art modeling procedures relative to this very simplistic baseline are minimal, and that the RMSEs that we obtain for these methods are almost as high as the standard deviations in the individual test sets (corresponding to the RMSE of the mean predictor).

## Discussion

The results of this paper are somewhat sobering: the performance of melting temperature prediction is substantially lower than what the current literature suggests, and current approaches to task-specific learning and fine-tuning struggle to learn features that generalize across species. There are a few caveats to these conclusions. First, the high Spearman rhos reported in the literature do reflect an ability to predict the effect on melting temperature on the overall amino acid composition. While they are not sensitive to the effect of individual mutations, we might reasonable expect that there is sufficient signal for e.g. de novo design protocols to design towards specific melting temperature profiles. Secondly, while we in this paper focus our attention on *supervised* learning strategies, it would be worth investigating how unsupervised models fare on this problem, specifically in light of the high performances reported on absolute free energy prediction Cagiada et al. [2024]. Finally, while we have tested a broad selection of methods, we have by no means exhaustively explored the space of modeling strategies. In particular, it is likely that better ways exist to incorporate 3D structure into the models. The log-likelihoods of inverse-folding models are known to correlate well with protein stability Boomsma and Frellsen [2017], Notin et al. [2023], and fine-tuning such models might therefore prove to generalize better across species.

The trade-off between predicting global and local properties of proteins is presumably not unique to melting temperature prediction. In the literature on variant effects, a distinction is sometimes made between global and local epistasis, where the former describes the genetic background in which variants occur, influencing the effects of observed point mutants Starr and Thornton [2016]. From this perspective, the cross-species models in our melting temperature experiments display an inclination towards capturing the diffuse effects of the genetic background rather than the specific effects of individual mutations. We might expect to see similar effects when predicting other protein properties, in particular those tied to specific/niche habitat conditions such as temperature, pH or salinity. When designing predictors for such tasks, it is therefore important that we design our metrics and train/validation/test setups to faithfully reflect the real-world application of interest. A notable positive example is found in the prediction of clinical variant effects in human populations, where substantial work has been done to document biases and temper overly optimistic performance estimates Livesey and Marsh [2020].

## Materials and Methods

### Embedding types

Our model employs two different embedding types: a sequence-based embedding from the ESM2 (Evolutionary Scale Modeling) model Lin et al. [2022] and an embedding obtained from the inverse folding model PiFold Hsu et al. [2022]. The former is an unsupervised language model based on the transformer architecture, trained on 250 million sequences from the UniRef dataset, which has been shown to capture and learn relevant information such as biochemical properties of amino acids, residue contact mapping, and homology detection, making it well-suited for various downstream tasks Rives et al. [2021], Lin et al. [2022], Michael et al. [2024]. As an inverse folding method, PiFold aims to predict protein sequences based on the atomic coordinates of protein backbone structures, and thus more directly captures the structural environment in its embeddings.

### Model design

The fundamental structure of the models is illustrated in Figure 1. The sequence and structural embeddings produce per-residue vector representations of 1280 and 128 dimensions respectively. These were either used independently or concatenated per residue and passed through a light attention block (see Figure 1). As the name suggests, light attention is a light-weight mechanism to approximate a traditional attention mechanism, and is commonly used as an alternative to simpler aggregation options such as calculating a mean over the sequence length Stärk et al. [2021]. This approach allows a unified and efficient representation of protein features, regardless of sequence length.

The light attention block was implemented using the default parameters as described in Stärk et al. [2021], although dimensionality of the input embeddings was adjusted based on the type of embedding used. For prediction, a multilayer perceptron (MLP) was used, comprising three hidden layers with an architecture of (128, 64, n), where n represents the number of output components, which varies depending on the specific experiment. This configuration allows for model adaptability to different predictive tasks while maintaining a consistent base structure.

### Rank-N-Contrast

The Rank-N-Contrast Zha et al. [2024] procedure consists of 3 components:

#### Data augmentation

A batch of input-label pairs undergoes data augmentation in the original embedding space using a Gaussian perturbation. While the original description of the method proposed constructing a two-view batch consisting of two types of augmentations, we deviate slightly from this by creating a two-view batch which comprises the original input and a transformed version of the same input.

#### Rank-N-Contrast Loss

A regression-aware loss function is constructed to learn a representation of the input data such that relative distances reflect differences in their continuous target values (i.e. melting temperatures). Following the original publication, the R*N* C loss per sample is defined as:

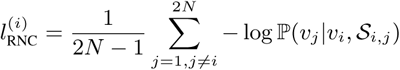

where:

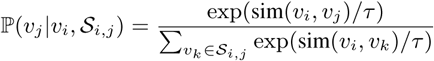

Here, sim(*·, ·*) is a similarity in representation-space between samples *vi* and *vj*, and *τ* is a temperature parameter. P(*vj |vi, Si,j*) represents the likelihood of a sample *vj* in the context of anchor *vi*, and the set of samples *Si,j*) that relative to *vi* have a worse rank than *vj* in terms of the *output* values.

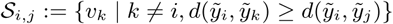

As an example, if *vj* is the sample that is closest to *vi* in terms of melting temperature, the likelihood would be optimized when *vj* had the highest similarity to *vi* in representation space. The full likelihood is simply an average over all samples in the batch.

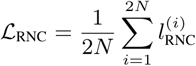

#### Representation Learning

The model learns an encoder by optimizing *L*RNC. Once the representation is learned, the encoder is frozen, and a separate predictor is trained using a standard loss. In our case, the learned Rank-N-Contrast representations are used as direct input to the light attention block in combination with MLP block in Figure 1.

### Data

All analyses were done on the Meltome partition of the FLIP dataset Dallago et al. [2021]. The dataset comprises 15 partitions designed to address various tasks in the field of protein design. For this study, we specifically focused on the partition related to thermostability, based on proteins with their corresponding melting temperatures (*Tm*) derived from the Meltome Atlas database. The ’Mix’ partition was selected, characterized by a high diversity of proteins originating from multiple species. Proteins less than 50 residues in length were discarded for this work. The amount of proteins corresponding to each species in the splits of the FLIP dataset is presented in the Table 2

**Table 2:**
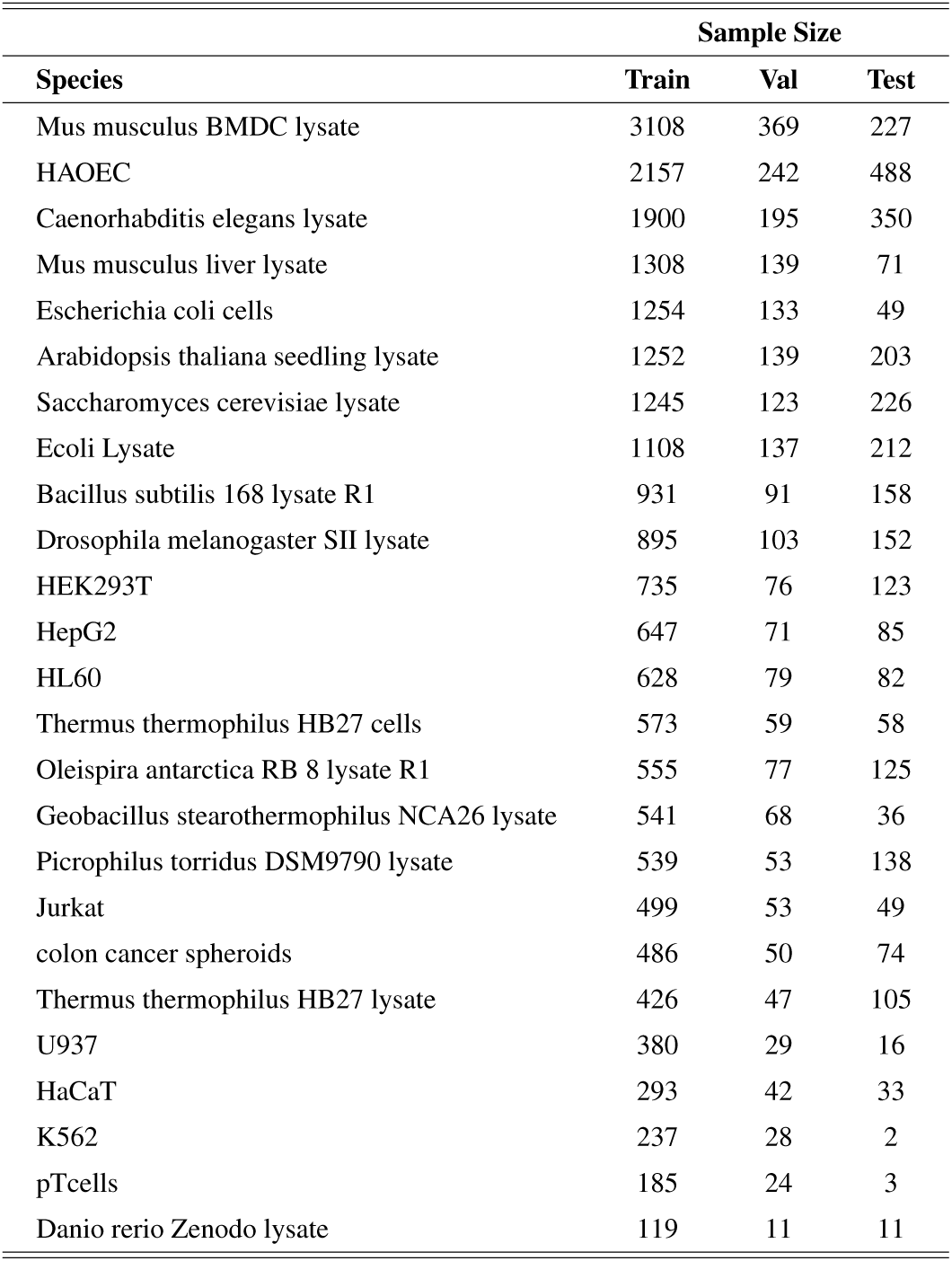
Sample distribution table by species. This table was constructed based on the distribution provided by the flip partition in relation to the Meltome Atlas, using the crossspecies partition or “mix”.

Protein sequences from the FLIP splits were used to compute embeddings using ESM2, which were then stored in files for efficient loading during model induction/inference. To map structural features, the UniProtIDs of each protein were used as a query in AlphaFoldDB Varadi et al. [2022] to obtain predicted structures. This approach was adopted to ensure the use of the closest predicted protein structure for each sequence, while avoiding the complexity associated with multi-chain protein structures. Once the protein structures were obtained, PiFold was used to generate graph embeddings, which were then stored in files. Notably, only the encoder component of the algorithm was used.

### Training

Model parameters were optimized using AdamW. The learning rate (lr) was adjusted according to the type of experiment: for the calibration of models using the full FLIP training partition in the global model (all species in the dataset), lr = 1 *×* 10^−4^ was set, while for the training of species-specific models (individual models trained with proteins belonging to the same species), lr = 1 *×* 10^−3^ was used.

Due to its particular architecture, the Rank N Contrast model followed a two-stage procedure, first learning representations using a contrastive procedure and then learning a regressor from these representations. The two components were optimised with different learning rates, denoted as lr*e* (encoder) and lr*p* (decoder). For global model calibration, lr*e* = 1 *×* 10^−5^ and lr*p* = 1 *×* 10^−4^, while for species-specific models, lr*e* = 1 *×* 10^−5^ and lr*p* = 1 *×* 10^−3^.

The fine-tuning approach was run exact as originally described Schmirler et al. (2024), following the provided Jupyter notebooks.

## Author contributions

The experimental design, along with the theoretical and methodological frameworks employed in this work, was developed collaboratively by all authors. SGL handled the algorithmic implementation, while the mathematical and conceptual formulations were jointly crafted by SGL and WB, with ideas refined iteratively to shape the final manuscript.

## Acknowledgments

This project has received funding from the European Union’s Horizon 2020 research and innovation programme under the Marie Sklodowska-Curie grant agreement No 801199, the Novo Nordisk Foundation through the MLSS Center (Basic Machine Learning Research in Life Science, NNF20OC0062606), and the Pioneer Centre for AI (DNRF grant number P1). The project was carried out in collaboration with the Enzyme Research Division of Novonesis A/S, located in Kongens Lyngby, Denmark.

## Financial disclosure

None reported.

## Conflict of interest

The authors declare no potential conflict of interests.

## Supporting information

Supplementary information, including comprehensive metric evaluations for all strategies used in this study and additional figures that further support the results, is provided in *Supplementary Material SA-SE*. These materials are available with the online version of the article.

## Suplementary Material

The supplementary material presented below includes supporting results for all experiments conducted in this study. This material includes violin plots and box plots for all methods and performance measures analysed in the paper, as well as tables with the corresponding results from the cross-species analysis.

### A Violin and Box Plots of Predictions Across Species

**Figure 6:**
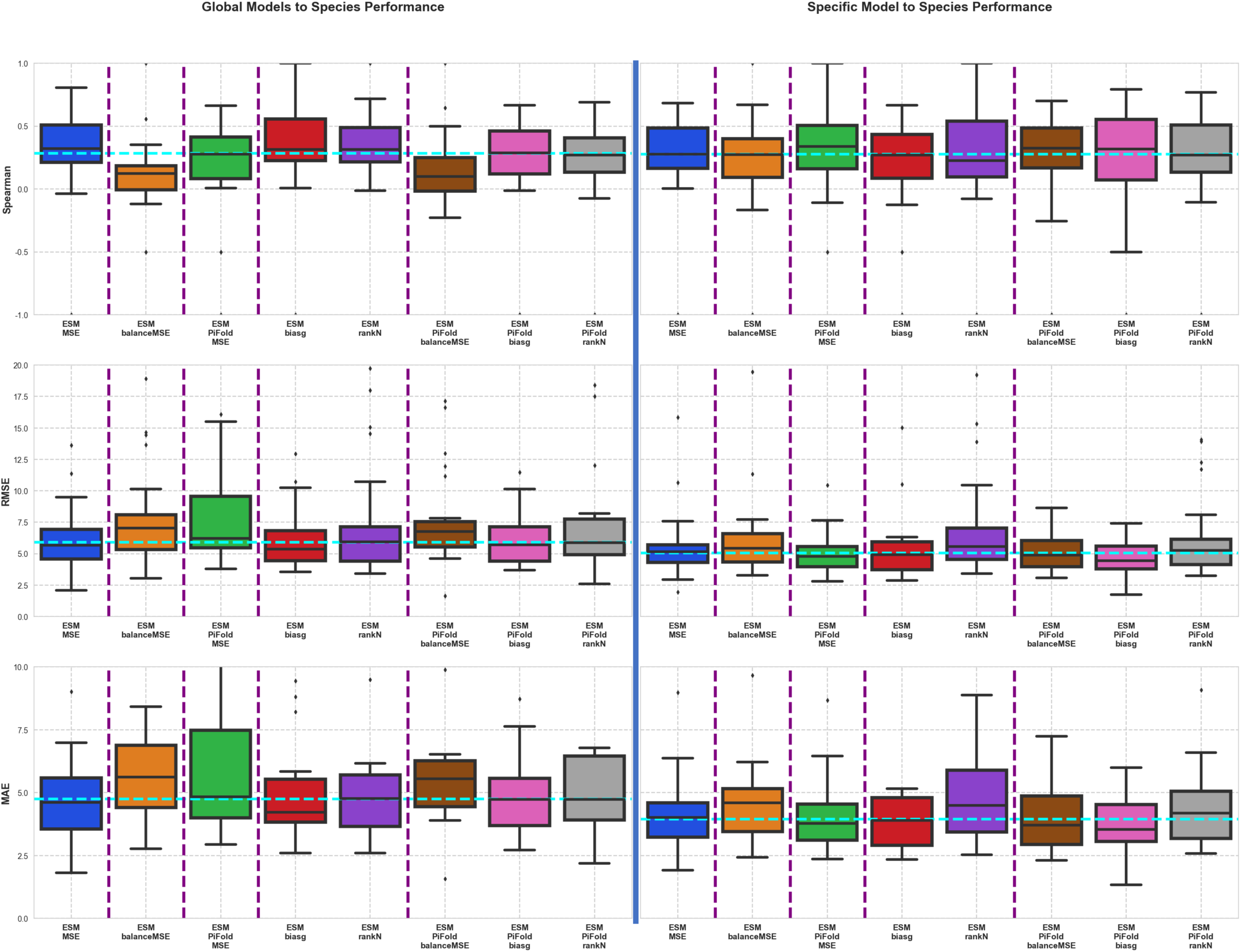
Box plots of the strategies used to predict thermostability. The blue line separates the use of global models applied to each species (right side) from the use of individual models for each species (left side). In addition, each region has subdivisions marked in purple, indicating the order in which the ideas were first tested: First, the prediction method using sequence embeddings was implemented. Next, species-by-species sampling was used in batches. Then the combination of sequence embeddings with inverse folding was used. Next, contrastive methods using sequence embeddings alone were applied. Finally, sequence embeddings were combined with contrastive methods. The cyan line crossing the box plots represents the average of the medians of all box plots. The points at the top and bottom of the box plots correspond to the outliers present in the predictions. Compared to the Boxplot shown in the main paper, this presents the boxplots related to all the performance metrics considered along the work, including: Spearman Correlation, RMSE and MAE respectively.

**Figure 7:**
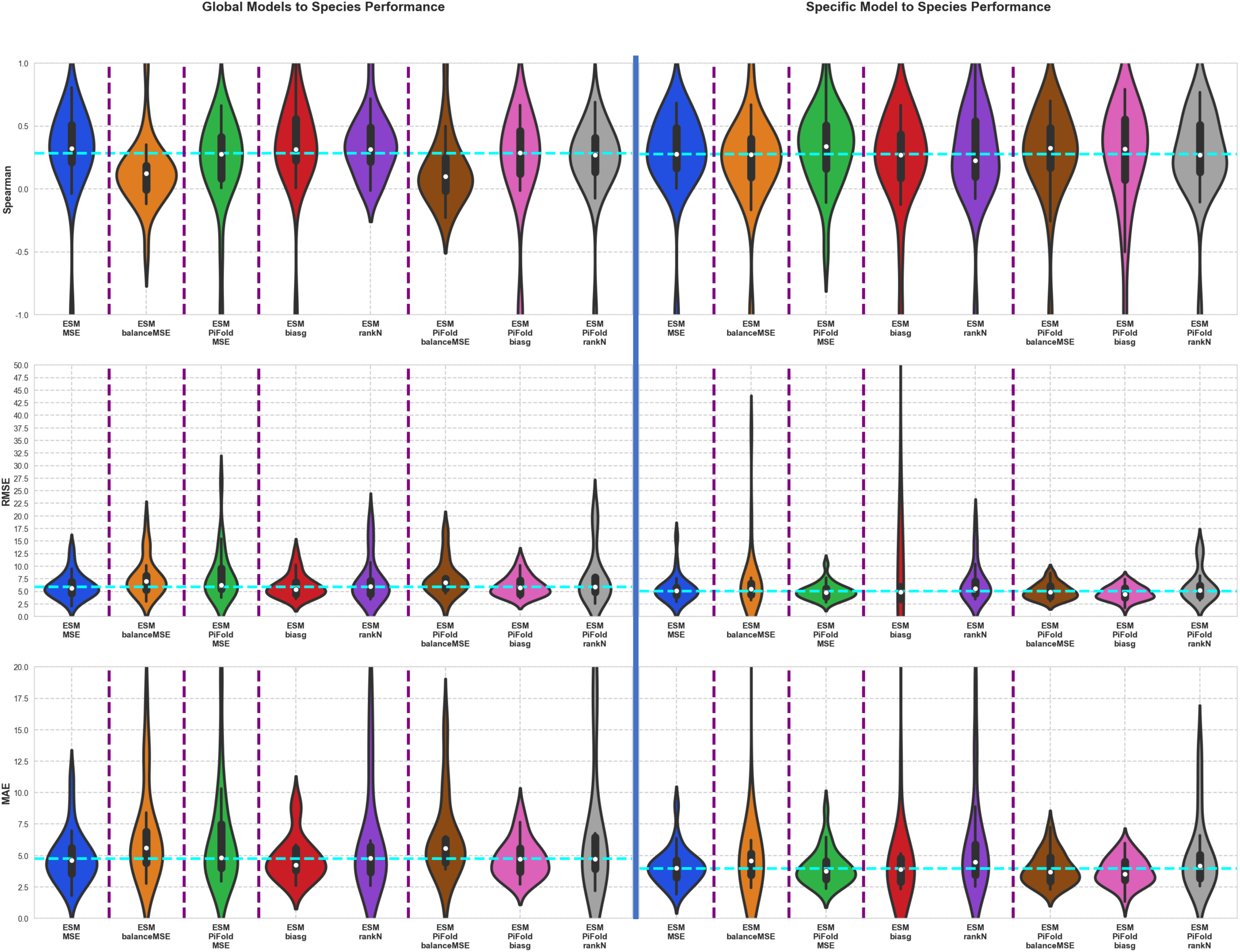
Violin plots of the strategies used to predict thermostability. The blue line separates the use of global models applied to each species (right side) from the use of individual models for each species (left side). In addition, each region has subdivisions marked in purple, indicating the order in which the ideas were first tested: First, the prediction method using sequence embeddings was implemented. Next, species-by-species sampling was used in batches. Then the combination of sequence embeddings with inverse folding was used. Next, contrastive methods using sequence embeddings alone were applied. Finally, sequence embeddings were combined with contrastive methods. The cyan line crossing the box plots represents the average of the medians of all violin plots. This plot show the variance and density estimation to all the performance metrics considered along the work, including: Spearman Correlation, RMSE and MAE respectively.

### B Tables of Results for the Full Flip Split

**Table 3:** Results of methods using sequence-based embeddings (ESM2). Performance evaluation using the full dataset of FLIP splits related to thermostability.

| Methods | global |  |  |  |
| --- | --- | --- | --- | --- |
|  | Spearman | MSE | RMSE | MAE |
| MSE | 0.688 | 37.303 | 6.1076 | 4.5268 |
| balanceMSE | 0.415 | 61.8778 | 7.8662 | 6.0651 |
| biasg | 0.6951 | 37.8031 | 6.1484 | 4.492 |
| rank_l2_l1_MSE_001_t2 | 0.6356 | 53.1513 | 7.2905 | 5.3756 |

**Table 4:** Results of methods using embeddings from protein structures using Inverse Folding Algorithms (PiFold). Performance evaluation using the full dataset of FLIP splits related to thermostability.

| Methods | global |  |  |  |
| --- | --- | --- | --- | --- |
|  | Spearman | MSE | RMSE | MAE |
| MSE | 0.3763 | 87.9824 | 9.3799 | 6.788 |
| balanceMSE | 0.3563 | 100.758 | 10.0378 | 7.1735 |
| biasg | 0.3961 | 88.4589 | 9.4053 | 6.6275 |
| rank_l2_l1_MSE_001_t2 | 0.309 | 98.5775 | 9.9286 | 7.0107 |

**Table 5:**
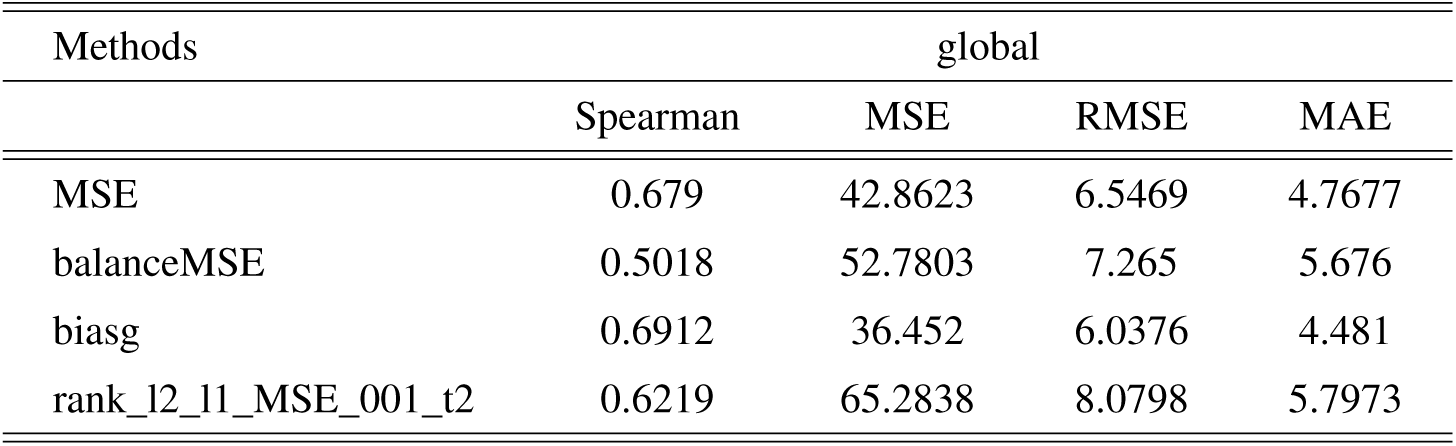
Results of methods using the concatenation of embeddings from ESM2 and PiFold. Performance evaluation using the full dataset of FLIP splits related to thermostability.

### C Tables of Results for Global Models Applied to Individual Species

**Table 6:**
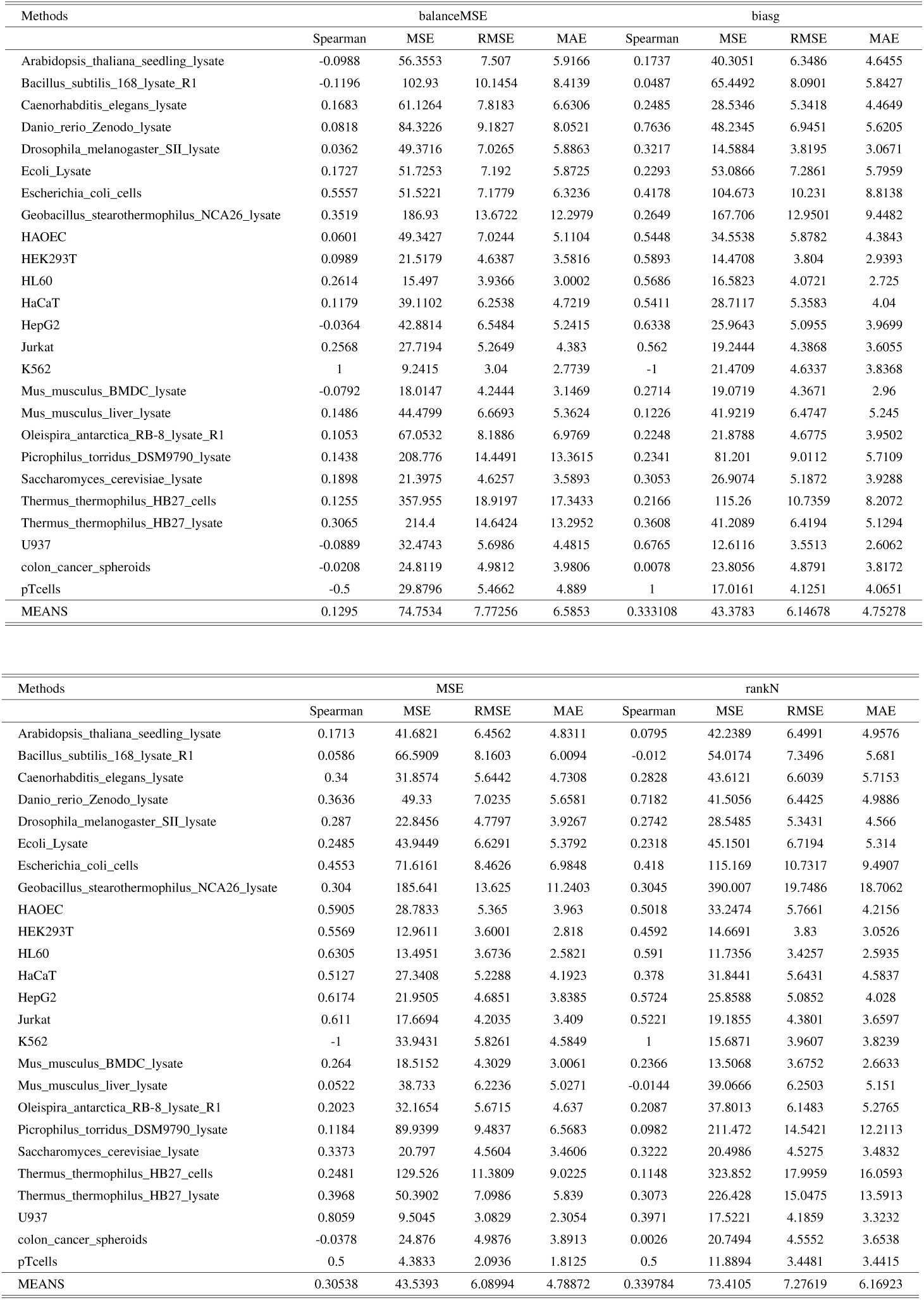
Evaluation of methods using sequence-based embeddings (ESM2): Performance evaluation was performed using the full training and validation sets of FLIP partitions related to thermostability to induce the model. Testing was performed on individual species within the FLIP test set. The table shows the evaluation based on four primary metrics: Spearman correlation, MSE, RMSE and MAE, for the four methods analysed in this study (balanceMSE, biasg, MSE and RankN).

| Methods | balanceMSE |  |  |  | biasg |  |  |  |
| --- | --- | --- | --- | --- | --- | --- | --- | --- |
|  | Spearman | MSE | RMSE | MAE | Spearman | MSE | RMSE | MAE |
| Arabidopsis_thaliana_seedling_lystate | -0.0988 | 56.3553 | 7.507 | 5.9166 | 0.1737 | 40.3051 | 6.3486 | 4.6455 |
| Bacillus_subtilis_168_lystate_R1 | -0.1196 | 102.93 | 10.1454 | 8.4139 | 0.0487 | 65.4492 | 8.0901 | 5.8427 |
| Caenorhabditis_elegans_lystate | 0.1683 | 61.1264 | 7.8183 | 6.6306 | 0.2485 | 28.5346 | 5.3418 | 4.4649 |
| Danio_rerio_Zenodo_lystate | 0.0818 | 84.3226 | 9.1827 | 8.0521 | 0.7636 | 48.2345 | 6.9451 | 5.6205 |
| Drosophila_melanogaster_SII_lystate | 0.0362 | 49.3716 | 7.0265 | 5.8863 | 0.3217 | 14.5884 | 3.8195 | 3.0671 |
| Ecoli_Lystate | 0.1727 | 51.7253 | 7.192 | 5.8725 | 0.2293 | 53.0866 | 7.2861 | 5.7959 |
| Escherichia_coli_cells | 0.5557 | 51.5221 | 7.1779 | 6.3236 | 0.4178 | 104.673 | 10.231 | 8.8138 |
| Geobacillus_stearothermophilus_NCA26_lystate | 0.3519 | 186.93 | 13.6722 | 12.2979 | 0.2649 | 167.706 | 12.9501 | 9.4482 |
| HAOEC | 0.0601 | 49.3427 | 7.0244 | 5.1104 | 0.5448 | 34.5538 | 5.8782 | 4.3843 |
| HEK293T | 0.0989 | 21.5179 | 4.6387 | 3.5816 | 0.5893 | 14.4708 | 3.804 | 2.9393 |
| HL60 | 0.2614 | 15.497 | 3.9366 | 3.0002 | 0.5686 | 16.5823 | 4.0721 | 2.725 |
| HaCaT | 0.1179 | 39.1102 | 6.2538 | 4.7219 | 0.5411 | 28.7117 | 5.3583 | 4.04 |
| HepG2 | -0.0364 | 42.8814 | 6.5484 | 5.2415 | 0.6338 | 25.9643 | 5.0955 | 3.9699 |
| Jurkat | 0.2568 | 27.7194 | 5.2649 | 4.383 | 0.562 | 19.2444 | 4.3868 | 3.6055 |
| K562 | 1 | 9.2415 | 3.04 | 2.7739 | -1 | 21.4709 | 4.6337 | 3.8368 |
| Mus_musculus_BMDC_lystate | -0.0792 | 18.0147 | 4.2444 | 3.1469 | 0.2714 | 19.0719 | 4.3671 | 2.96 |
| Mus_musculus_liver_lystate | 0.1486 | 44.4799 | 6.6693 | 5.3624 | 0.1226 | 41.9219 | 6.4747 | 5.245 |
| Oleispira_antarctica_RB-8_lystate_R1 | 0.1053 | 67.0532 | 8.1886 | 6.9769 | 0.2248 | 21.8788 | 4.6775 | 3.9502 |
| Picrophilus_torridus_DSM9790_lystate | 0.1438 | 208.776 | 14.4491 | 13.3615 | 0.2341 | 81.201 | 9.0112 | 5.7109 |
| Saccharomyces_cerevisiae_lystate | 0.1898 | 21.3975 | 4.6257 | 3.5893 | 0.3053 | 26.9074 | 5.1872 | 3.9288 |
| Thermus_thermophilus_HB27_cells | 0.1255 | 357.955 | 18.9197 | 17.3433 | 0.2166 | 115.26 | 10.7359 | 8.2072 |
| Thermus_thermophilus_HB27_lystate | 0.3065 | 214.4 | 14.6424 | 13.2952 | 0.3608 | 41.2089 | 6.4194 | 5.1294 |
| U937 | -0.0889 | 32.4743 | 5.6986 | 4.4815 | 0.6765 | 12.6116 | 3.5513 | 2.6062 |
| colon_cancer_spheroids | -0.0208 | 24.8119 | 4.9812 | 3.9806 | 0.0078 | 23.8056 | 4.8791 | 3.8172 |
| pTcells | -0.5 | 29.8796 | 5.4662 | 4.889 | 1 | 17.0161 | 4.1251 | 4.0651 |
| MEANS | 0.1295 | 74.7534 | 7.77256 | 6.5853 | 0.333108 | 43.3783 | 6.14678 | 4.75278 |

Table 6: Evaluation of methods using sequence-based embeddings (ESM2): Performance evaluation was performed using the full training and validation sets of FLIP partitions related to thermostability to induce the model.
| Methods | MSE |  |  |  | rankN |  |  |  |
| --- | --- | --- | --- | --- | --- | --- | --- | --- |
|  | Spearman | MSE | RMSE | MAE | Spearman | MSE | RMSE | MAE |
| Arabidopsis_thaliana_seedling_lystate | 0.1713 | 41.6821 | 6.4562 | 4.8311 | 0.0795 | 42.2389 | 6.4991 | 4.9576 |
| Bacillus_subtilis_168_lystate_R1 | 0.0586 | 66.5909 | 8.1603 | 6.0094 | -0.012 | 54.0174 | 7.3496 | 5.681 |
| Caenorhabditis_elegans_lystate | 0.34 | 31.8574 | 5.6442 | 4.7308 | 0.2828 | 43.6121 | 6.6039 | 5.7153 |
| Danio_rerio_Zenodo_lystate | 0.3636 | 49.33 | 7.0235 | 5.6581 | 0.7182 | 41.5056 | 6.4425 | 4.9886 |
| Drosophila_melanogaster_SII_lystate | 0.287 | 22.8456 | 4.7797 | 3.9267 | 0.2742 | 28.5485 | 5.3431 | 4.566 |
| Ecoli_Lystate | 0.2485 | 43.9449 | 6.6291 | 5.3792 | 0.2318 | 45.1501 | 6.7194 | 5.314 |
| Escherichia_coli_cells | 0.4553 | 71.6161 | 8.4626 | 6.9848 | 0.418 | 115.169 | 10.7317 | 9.4907 |
| Geobacillus_stearothermophilus_NCA26_lystate | 0.304 | 185.641 | 13.625 | 11.2403 | 0.3045 | 390.007 | 19.7486 | 18.7062 |
| HAOEC | 0.5905 | 28.7833 | 5.365 | 3.963 | 0.5018 | 33.2474 | 5.7661 | 4.2156 |
| HEK293T | 0.5569 | 12.9611 | 3.6001 | 2.818 | 0.4592 | 14.6691 | 3.83 | 3.0526 |
| HL60 | 0.6305 | 13.4951 | 3.6736 | 2.5821 | 0.591 | 11.7356 | 3.4257 | 2.5935 |
| HaCaT | 0.5127 | 27.3408 | 5.2288 | 4.1923 | 0.378 | 31.8441 | 5.6431 | 4.5837 |
| HepG2 | 0.6174 | 21.9505 | 4.6851 | 3.8385 | 0.5724 | 25.8588 | 5.0852 | 4.028 |
| Jurkat | 0.611 | 17.6694 | 4.2035 | 3.409 | 0.5221 | 19.1855 | 4.3801 | 3.6597 |
| K562 | -1 | 33.9431 | 5.8261 | 4.5849 | 1 | 15.6871 | 3.9607 | 3.8239 |
| Mus_musculus_BMDC_lystate | 0.264 | 18.5152 | 4.3029 | 3.0061 | 0.2366 | 13.5068 | 3.6752 | 2.6633 |
| Mus_musculus_liver_lystate | 0.0522 | 38.733 | 6.2236 | 5.0271 | -0.0144 | 39.0666 | 6.2503 | 5.151 |
| Oleispira_antarctica_RB-8_lystate_R1 | 0.2023 | 32.1654 | 5.6715 | 4.637 | 0.2087 | 37.8013 | 6.1483 | 5.2765 |
| Picrophilus_torridus_DSM9790_lystate | 0.1184 | 89.9399 | 9.4837 | 6.5683 | 0.0982 | 211.472 | 14.5421 | 12.2113 |
| Saccharomyces_cerevisiae_lystate | 0.3373 | 20.797 | 4.5604 | 3.4606 | 0.3222 | 20.4986 | 4.5275 | 3.4832 |
| Thermus_thermophilus_HB27_cells | 0.2481 | 129.526 | 11.3809 | 9.0225 | 0.1148 | 323.852 | 17.9959 | 16.0593 |
| Thermus_thermophilus_HB27_lystate | 0.3968 | 50.3902 | 7.0986 | 5.839 | 0.3073 | 226.428 | 15.0475 | 13.5913 |
| U937 | 0.8059 | 9.5045 | 3.0829 | 2.3054 | 0.3971 | 17.5221 | 4.1859 | 3.3232 |
| colon_cancer_spheroids | -0.0378 | 24.876 | 4.9876 | 3.8913 | 0.0026 | 20.7494 | 4.5552 | 3.6538 |
| pTcells | 0.5 | 4.3833 | 2.0936 | 1.8125 | 0.5 | 11.8894 | 3.4481 | 3.4415 |
| MEANS | 0.30538 | 43.5393 | 6.08994 | 4.78872 | 0.339784 | 73.4105 | 7.27619 | 6.16923 |

**Table 7:** Evaluation of methods using embeddings from protein structures via inverse folding (PiFold): Performance evaluation was performed using the full training and validation sets of FLIP partitions related to thermostability to induce the model. Testing was performed on individual species within the FLIP test set. The table shows the evaluation based on four primary metrics: Spearman correlation, MSE, RMSE and MAE, for the four methods analysed in this study (balanceMSE, biasg, MSE and RankN).

| Methods | balanceMSE |  |  |  | biasg |  |  |  |
| --- | --- | --- | --- | --- | --- | --- | --- | --- |
|  | Spearman | MSE | RMSE | MAE | Spearman | MSE | RMSE | MAE |
| Arabidopsis_thaliana_seedling_lystate | 0.1978 | 43.6081 | 6.6036 | 5.1771 | 0.0682 | 47.5744 | 6.8974 | 5.3945 |
| Bacillus_subtilis_168_lystate_R1 | 0.1186 | 64.2171 | 8.0136 | 6.2432 | 0.1013 | 90.2633 | 9.5007 | 7.0436 |
| Caenorhabditis_elegans_lystate | 0.1135 | 77.9945 | 8.8314 | 7.4103 | 0.1488 | 67.6063 | 8.2223 | 6.8556 |
| Danio_rerio_Zenodo_lystate | 0.1545 | 67.2587 | 8.2011 | 5.6998 | 0.4818 | 85.5648 | 9.2501 | 7.8557 |
| Drosophila_melanogaster_SII_lystate | 0.1883 | 42.6002 | 6.5269 | 5.436 | 0.0446 | 51.1427 | 7.1514 | 5.9902 |
| Ecoli_lystate | 0.0712 | 61.3955 | 7.8355 | 6.194 | 0.101 | 72.936 | 8.5403 | 6.6497 |
| Escherichia_coli_cells | 0.2221 | 135.125 | 11.6243 | 10.0279 | 0.0373 | 176.004 | 13.2666 | 11.8437 |
| Geobacillus_stearothermophilus_NCA26_lystate | 0.088 | 498.251 | 22.3215 | 21.2174 | 0.0049 | 478.348 | 21.8712 | 19.6788 |
| HAOEC | 0.3888 | 49.0695 | 7.005 | 5.4388 | 0.3905 | 41.4828 | 6.4407 | 4.8899 |
| HEK293T | 0.3737 | 24.4279 | 4.9425 | 3.768 | 0.3315 | 21.264 | 4.6113 | 3.5963 |
| HL60 | 0.2711 | 23.8911 | 4.8879 | 3.6236 | 0.3725 | 17.2556 | 4.154 | 3.2397 |
| HaCaT | 0.2216 | 45.1755 | 6.7213 | 4.7603 | 0.242 | 37.8045 | 6.1485 | 4.8963 |
| HepG2 | 0.3273 | 38.7255 | 6.223 | 4.6599 | 0.4269 | 34.835 | 5.9021 | 4.8028 |
| Jurkat | 0.1976 | 34.0146 | 5.8322 | 4.6808 | 0.4273 | 25.7759 | 5.077 | 4.1718 |
| K562 | -1 | 4.0299 | 2.0075 | 1.5151 | -1 | 18.3807 | 4.2873 | 4.287 |
| Mus_musculus_BMDC_lystate | 0.1254 | 26.6945 | 5.1667 | 3.6526 | 0.1506 | 27.9881 | 5.2904 | 3.6731 |
| Mus_musculus_liver_lystate | 0.1326 | 46.4017 | 6.8119 | 5.5285 | 0.1947 | 46.4456 | 6.8151 | 5.6925 |
| Oleispira_antartica_RB-8_lystate_R1 | 0.1121 | 59.6047 | 7.7204 | 6.3074 | 0.0632 | 61.3767 | 7.8343 | 6.0074 |
| Picrophilus_torridus_DSM9790_lystate | -0.041 | 410.771 | 20.2675 | 18.9768 | 0.126 | 305.723 | 17.4849 | 14.515 |
| Saccharomyces_cerevisiae_lystate | 0.053 | 32.4263 | 5.6944 | 4.5244 | 0.3205 | 29.4371 | 5.4256 | 4.2492 |
| Thermus_thermophilus_HB27_cells | -0.1055 | 838.375 | 28.9547 | 26.1275 | -0.0737 | 642.825 | 25.354 | 20.9251 |
| Thermus_thermophilus_HB27_lystate | 0.0468 | 582.313 | 24.1312 | 21.609 | -0.0441 | 483.995 | 21.9999 | 17.1478 |
| U937 | 0.3706 | 25.0477 | 5.0048 | 4.2869 | 0.1 | 30.4037 | 5.514 | 4.1933 |
| colon_cancer_spheroids | 0.0613 | 26.1402 | 5.1127 | 4.1275 | 0.2479 | 38.4994 | 6.2048 | 4.5201 |
| pTcells | 0.5 | 34.794 | 5.8986 | 5.8971 | -0.5 | 50.3598 | 7.0965 | 6.875 |
| MEANS | 0.127576 | 131.694 | 9.29361 | 7.8756 | 0.110548 | 119.332 | 9.21362 | 7.55976 |

Table 7: Evaluation of methods using embeddings from protein structures via inverse folding (PiFold): Performance evaluation was performed using the full training and validation sets of FLIP partitions related to thermostability to induce the model.
| Methods | MSE |  |  |  | rankN |  |  |  |
| --- | --- | --- | --- | --- | --- | --- | --- | --- |
|  | Spearman | MSE | RMSE | MAE | Spearman | MSE | RMSE | MAE |
| Arabidopsis_thaliana_seedling_lystate | 0.0615 | 49.9119 | 7.0648 | 5.4569 | 0.04 | 47.601 | 6.8993 | 5.3863 |
| Bacillus_subtilis_168_lystate_R1 | 0.0577 | 77.1768 | 8.785 | 6.8443 | -0.0649 | 67.5899 | 8.2213 | 6.6734 |
| Caenorhabditis_elegans_lystate | 0.2483 | 88.5995 | 9.4127 | 8.0938 | -0.0366 | 76.9103 | 8.7699 | 7.6532 |
| Danio_rerio_Zenodo_lystate | -0.0182 | 63.064 | 7.9413 | 5.9462 | 0.6545 | 53.9727 | 7.3466 | 5.702 |
| Drosophila_melanogaster_SII_lystate | 0.2405 | 56.1757 | 7.495 | 6.493 | 0.3073 | 40.6112 | 6.3727 | 5.6224 |
| Ecoli_lystate | 0.0873 | 66.5738 | 8.1593 | 6.4321 | -0.0549 | 59.2102 | 7.6948 | 6.0148 |
| Escherichia_coli_cells | 0.0935 | 141.508 | 11.8957 | 10.15 | 0.0313 | 108.28 | 10.4057 | 9.1288 |
| Geobacillus_stearothermophilus_NCA26_lystate | -0.0762 | 402.561 | 20.0639 | 18.2502 | -0.2718 | 397.585 | 19.9395 | 18.0143 |
| HAOEC | 0.389 | 37.326 | 6.1095 | 4.579 | 0.3736 | 47.5098 | 6.8927 | 5.0855 |
| HEK293T | 0.3936 | 23.1623 | 4.8127 | 3.5236 | 0.3369 | 23.2582 | 4.8227 | 3.5999 |
| HL60 | 0.3305 | 19.1996 | 4.3817 | 3.4348 | 0.354 | 19.2183 | 4.3839 | 3.1957 |
| HaCaT | 0.0785 | 40.7701 | 6.3851 | 5.359 | 0.3414 | 39.0351 | 6.2478 | 4.8033 |
| HepG2 | 0.5175 | 25.7657 | 5.076 | 4.0447 | 0.4026 | 34.4372 | 5.8683 | 4.7732 |
| Jurkat | 0.5916 | 17.522 | 4.1859 | 3.4371 | 0.2134 | 31.7117 | 5.6313 | 4.5822 |
| K562 | 1 | 5.6946 | 2.3863 | 2.1568 | 0 | 5.059 | 2.2492 | 1.8709 |
| Mus_musculus_BMDC_lystate | 0.21 | 20.926 | 4.5745 | 3.4033 | 0.1203 | 16.1929 | 4.024 | 2.9002 |
| Mus_musculus_liver_lystate | -0.0108 | 46.809 | 6.8417 | 5.7137 | -0.0597 | 48.1737 | 6.9407 | 5.7201 |
| Oleispira_antartica_RB-8_lystate_R1 | 0.0712 | 74.8146 | 8.6495 | 7.3418 | 0.1333 | 72.2167 | 8.498 | 7.3424 |
| Picrophilus_torridus_DSM9790_lystate | 0.1537 | 314.616 | 17.7374 | 15.7521 | -0.0237 | 393.507 | 19.837 | 18.6257 |
| Saccharomyces_cerevisiae_lystate | 0.1838 | 21.5273 | 4.6398 | 3.6465 | 0.1058 | 21.5062 | 4.6375 | 3.6898 |
| Thermus_thermophilus_HB27_cells | -0.1581 | 815.522 | 28.5573 | 25.5008 | -0.013 | 860.323 | 29.3313 | 26.241 |
| Thermus_thermophilus_HB27_lystate | 0.0875 | 492.982 | 22.2032 | 19.2164 | 0.0335 | 609.46 | 24.6873 | 22.0347 |
| U937 | 0.4765 | 22.0209 | 4.6926 | 3.7639 | 0.3316 | 28.9023 | 5.3761 | 4.2117 |
| colon_cancer_spheroids | 0.3813 | 23.1553 | 4.812 | 3.6682 | 0.27 | 18.2909 | 4.2768 | 3.5334 |
| pTcells | 0.5 | 46.0152 | 6.7835 | 6.4249 | -0.5 | 36.0084 | 6.0007 | 5.9248 |
| MEANS | 0.235608 | 119.736 | 8.94586 | 7.54532 | 0.120996 | 126.263 | 9.0142 | 7.69319 |

**Table 8:** Evaluation of methods using concatenation of embeddings from ESM2 and PiFold: Performance evaluation was performed using the full training and validation sets of FLIP partitions related to thermostability to induce the model. Testing was performed on individual species within the FLIP test set. The table shows the evaluation based on four primary metrics: Spearman correlation, MSE, RMSE and MAE, for the four methods analysed in this study (balanceMSE, biasg, MSE and RankN).

| Methods | balanceMSE |  |  |  | biasg |  |  |  |
| --- | --- | --- | --- | --- | --- | --- | --- | --- |
|  | Spearman | MSE | RMSE | MAE | Spearman | MSE | RMSE | MAE |
| Arabidopsis_thaliana_seedling_lystate | -0.0328 | 44.9936 | 6.7077 | 5.1725 | 0.1137 | 42.7874 | 6.5412 | 5.0204 |
| Bacillus_subtilis_168_lystate_R1 | 0.0251 | 48.0577 | 6.9324 | 5.6365 | -0.0123 | 75.4061 | 8.6837 | 6.5255 |
| Caenorhabditis_elegans_lystate | 0.1034 | 45.9781 | 6.7807 | 5.8325 | 0.3285 | 32.324 | 5.6854 | 4.6984 |
| Danio_rerio_Zenodo_lystate | 0.6455 | 61.3115 | 7.8302 | 6.5264 | 0.6455 | 46.4755 | 6.8173 | 5.611 |
| Drosophila_melanogaster_SII_lystate | 0.1124 | 25.6896 | 5.0685 | 4.1189 | 0.3215 | 20.6017 | 4.5389 | 3.6341 |
| Ecoli_lystate | 0.0858 | 50.1222 | 7.0797 | 5.7697 | 0.2931 | 43.929 | 6.6279 | 5.4187 |
| Escherichia_coli_cells | 0.3961 | 124.554 | 11.1604 | 9.8918 | 0.3531 | 72.1184 | 8.4923 | 7.3403 |
| Geobacillus_stearothermophilus_NCA26_lystate | 0.2636 | 293.461 | 17.1307 | 15.6566 | 0.2785 | 131.69 | 11.4756 | 8.7325 |
| HAOEC | -0.0117 | 52.6367 | 7.2551 | 5.5859 | 0.5974 | 28.9445 | 5.38 | 3.9445 |
| HEK293T | -0.0166 | 30.0877 | 5.4852 | 4.3884 | 0.4751 | 16.0743 | 4.0093 | 3.0436 |
| HL60 | -0.026 | 27.6587 | 5.2592 | 4.0565 | 0.5124 | 13.6517 | 3.6948 | 2.713 |
| HaCaT | -0.2289 | 55.3317 | 7.4385 | 5.8553 | 0.6654 | 24.5686 | 4.9567 | 4.0428 |
| HepG2 | -0.0209 | 47.0074 | 6.8562 | 5.5277 | 0.5767 | 23.547 | 4.8525 | 3.8697 |
| Jurkat | -0.2288 | 45.9317 | 6.7773 | 5.9716 | 0.5776 | 16.0434 | 4.0054 | 3.3378 |
| K562 | 1 | 2.7174 | 1.6485 | 1.5705 | -1 | 55.7615 | 7.4674 | 5.3456 |
| Mus_musculus_BMDC_lystate | -0.0049 | 23.717 | 4.87 | 3.8992 | 0.2541 | 18.3161 | 4.2797 | 2.9783 |
| Mus_musculus_liver_lystate | 0.1471 | 43.8143 | 6.6192 | 5.4036 | 0.1062 | 38.0584 | 6.1692 | 4.8434 |
| Oleispira_antartica_RB-8_lystate_R1 | 0.1304 | 32.0531 | 5.6616 | 4.8655 | 0.1592 | 32.9704 | 5.742 | 4.7691 |
| Picrophilus_torridus_DSM9790_lystate | 0.0946 | 168.576 | 12.9837 | 10.7069 | -0.0145 | 80.3387 | 8.9632 | 6.2426 |
| Saccharomyces_cerevisiae_lystate | 0.0423 | 30.4126 | 5.5148 | 4.3921 | 0.3267 | 18.9499 | 4.3531 | 3.2769 |
| Thermus_thermophilus_HB27_cells | 0.2346 | 276.846 | 16.6387 | 14.5587 | 0.1349 | 102.891 | 10.1435 | 7.6395 |
| Thermus_thermophilus_HB27_lystate | 0.3538 | 143.249 | 11.9687 | 10.2945 | 0.2823 | 52.6987 | 7.2594 | 5.6441 |
| U937 | 0.2529 | 33.3963 | 5.779 | 4.7876 | 0.4147 | 14.9665 | 3.8687 | 3.0603 |
| colon_cancer_spheroids | -0.1639 | 31.1899 | 5.5848 | 4.6067 | 0.0389 | 24.9092 | 4.9909 | 3.8851 |
| pTcells | 0.5 | 21.1659 | 4.6006 | 4.2683 | -1 | 19.0944 | 4.3697 | 3.8958 |
| MEANS | 0.146124 | 70.3984 | 7.58526 | 6.37376 | 0.217148 | 41.8846 | 6.13471 | 4.78052 |

Table 8: Evaluation of methods using concatenation of embeddings from ESM2 and PiFold: Performance evaluation was performed using the full training and validation sets of FLIP partitions related to thermostability to induce the model.
| Methods | MSE |  |  |  | rankN |  |  |  |
| --- | --- | --- | --- | --- | --- | --- | --- | --- |
|  | Spearman | MSE | RMSE | MAE | Spearman | MSE | RMSE | MAE |
| Arabidopsis_thaliana_seedling_lystate | 0.109 | 39.2262 | 6.2631 | 4.8214 | 0.1274 | 36.2154 | 6.0179 | 4.5516 |
| Bacillus_subtilis_168_lystate_R1 | 0.0245 | 59.9784 | 7.7446 | 5.7714 | 0.0426 | 37.9457 | 6.16 | 4.8815 |
| Caenorhabditis_elegans_lystate | 0.3299 | 32.1148 | 5.667 | 4.8481 | 0.2677 | 33.5518 | 5.7924 | 4.8652 |
| Danio_rerio_Zenodo_lystate | 0.6636 | 47.1145 | 6.864 | 5.3625 | 0.6909 | 67.0075 | 8.1858 | 6.6925 |
| Drosophila_melanogaster_SII_lystate | 0.2991 | 22.454 | 4.7386 | 3.8411 | 0.381 | 17.4374 | 4.1758 | 3.4377 |
| Ecoli_lystate | 0.2094 | 47.4049 | 6.8851 | 5.5404 | 0.2948 | 52.1178 | 7.2193 | 5.7509 |
| Escherichia_coli_cells | 0.3589 | 99.9341 | 9.9967 | 8.4899 | 0.2228 | 144.307 | 12.0128 | 10.624 |
| Geobacillus_stearothermophilus_NCA26_lystate | 0.2144 | 258.657 | 16.0828 | 13.9394 | 0.271 | 464.474 | 21.5517 | 20.7134 |
| HAOEC | 0.5121 | 32.7875 | 5.726 | 4.1267 | 0.4289 | 45.9515 | 6.7788 | 5.0711 |
| HEK293T | 0.4918 | 14.5158 | 3.81 | 2.9418 | 0.3636 | 18.5763 | 4.31 | 3.312 |
| HL60 | 0.5576 | 25.8021 | 5.0796 | 3.0008 | 0.4153 | 15.7443 | 3.9679 | 2.8836 |
| HaCaT | 0.4221 | 29.4583 | 5.4275 | 4.4413 | 0.2096 | 35.3295 | 5.9439 | 4.6046 |
| HepG2 | 0.5828 | 31.2366 | 5.589 | 4.0038 | 0.5321 | 31.8896 | 5.6471 | 4.3951 |
| Jurkat | 0.5608 | 34.4767 | 5.8717 | 4.0023 | 0.426 | 25.9512 | 5.0942 | 4.2632 |
| K562 | -1 | 714.098 | 26.7226 | 19.1323 | -1 | 6.7103 | 2.5904 | 2.1838 |
| Mus_musculus_BMDC_lystate | 0.266 | 16.9931 | 4.1223 | 3.0564 | 0.2084 | 14.4295 | 3.7986 | 2.7389 |
| Mus_musculus_liver_lystate | 0.0062 | 40.8254 | 6.3895 | 5.0758 | 0.0941 | 48.9111 | 6.9936 | 5.7595 |
| Oleispira_antartica_RB-8_lystate_R1 | 0.1337 | 24.9606 | 4.9961 | 4.1284 | 0.1489 | 27.9463 | 5.2864 | 4.5097 |
| Picrophilus_torridus_DSM9790_lystate | 0.0185 | 119.918 | 10.9507 | 8.2336 | 0.0208 | 338.396 | 18.3955 | 16.5375 |
| Saccharomyces_cerevisiae_lystate | 0.3261 | 20.3761 | 4.514 | 3.4657 | 0.2844 | 27.5199 | 5.2459 | 4.2572 |
| Thermus_thermophilus_HB27_cells | 0.073 | 170.008 | 13.0387 | 10.0611 | 0.0676 | 431.125 | 20.7636 | 18.904 |
| Thermus_thermophilus_HB27_lystate | 0.2913 | 120.471 | 10.9759 | 7.906 | 0.2938 | 307.381 | 17.5323 | 16.1797 |
| U937 | 0.3941 | 38.377 | 6.1949 | 3.9725 | 0.6 | 19.7639 | 4.4457 | 3.4367 |
| colon_cancer_spheroids | 0.0681 | 33.1918 | 5.7612 | 4.0455 | -0.0761 | 23.4299 | 4.8404 | 3.8038 |
| pTcells | -0.5 | 239.848 | 15.487 | 10.3164 | 0.5 | 27.3404 | 5.2288 | 5.0902 |
| MEANS | 0.21652 | 92.5691 | 8.19594 | 6.18098 | 0.232624 | 91.9781 | 7.91915 | 6.7779 |

### D Tables of Results for Individual Model per Species

**Table 9:**
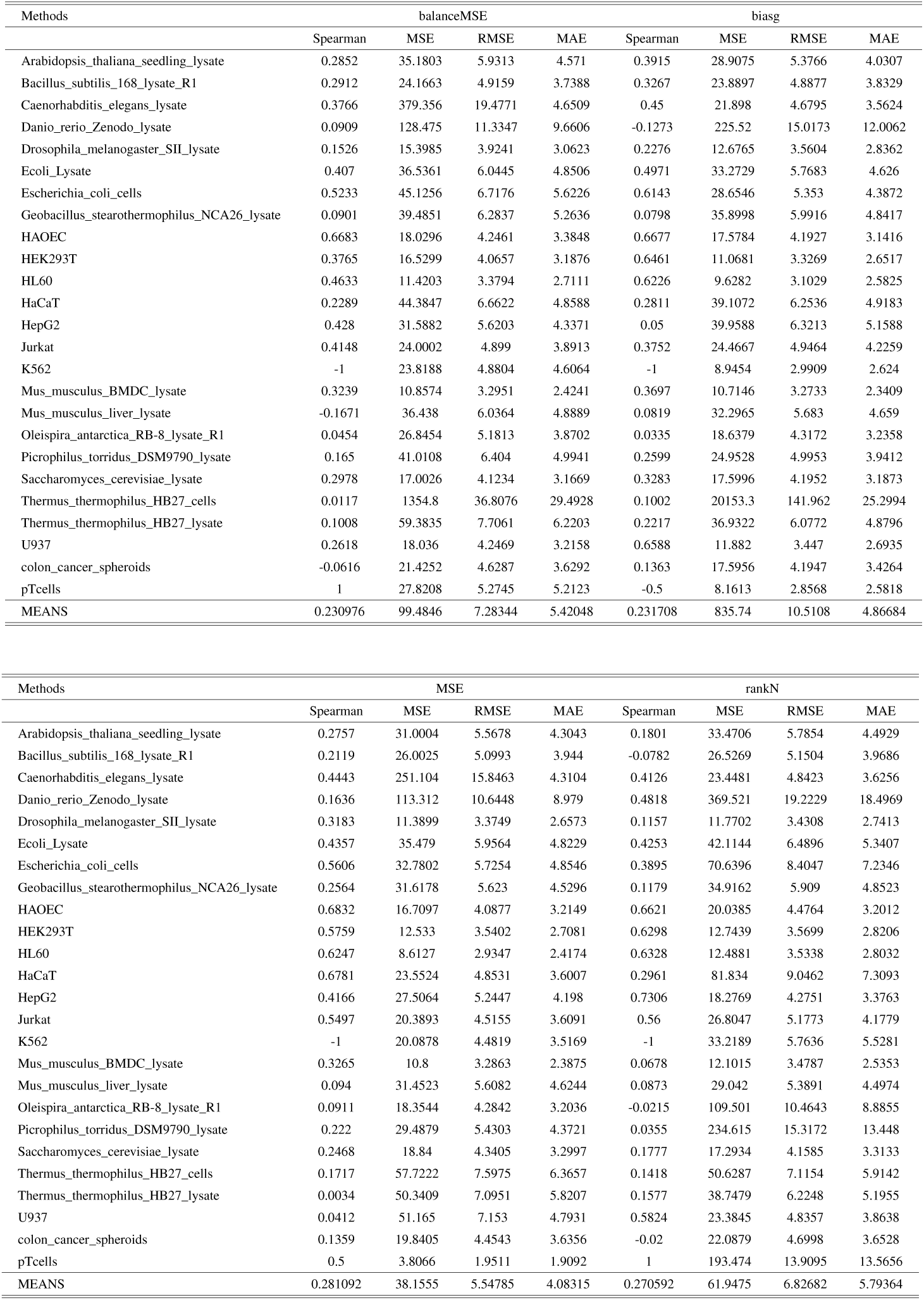
Assessment of methodologies employing sequence-based embeddings from ESM2: The performance of individual models was assessed for each species and subsequently tested on that target species inside the FLIP partition. Spearman correlation, mean square error (MSE), root mean square error (RMSE) and mean absolute error (MAE). These metrics were used to analyse the effectiveness of the four methods investigated in this study: balanceMSE, biasg, MSE and RankN.

| Methods | balanceMSE |  |  |  | biasg |  |  |  |
| --- | --- | --- | --- | --- | --- | --- | --- | --- |
|  | Spearman | MSE | RMSE | MAE | Spearman | MSE | RMSE | MAE |
| Arabidopsis_thaliana_seedling_lystate | 0.2852 | 35.1803 | 5.9313 | 4.571 | 0.3915 | 28.9075 | 5.3766 | 4.0307 |
| Bacillus_subtilis_168_lystate_R1 | 0.2912 | 24.1663 | 4.9159 | 3.7388 | 0.3267 | 23.8897 | 4.8877 | 3.8329 |
| Caenorhabditis_elegans_lystate | 0.3766 | 379.356 | 19.4771 | 4.6509 | 0.45 | 21.898 | 4.6795 | 3.5624 |
| Danio_rerio_Zenodo_lystate | 0.0909 | 128.475 | 11.3347 | 9.6606 | -0.1273 | 225.52 | 15.0173 | 12.0062 |
| Drosophila_melanogaster_SII_lystate | 0.1526 | 15.3985 | 3.9241 | 3.0623 | 0.2276 | 12.6765 | 3.5604 | 2.8362 |
| Ecoli_Lysate | 0.407 | 36.5361 | 6.0445 | 4.8506 | 0.4971 | 33.2729 | 5.7683 | 4.626 |
| Escherichia_coli_cells | 0.5233 | 45.1256 | 6.7176 | 5.6226 | 0.6143 | 28.6546 | 5.353 | 4.3872 |
| Geobacillus_stearothermophilus_NCA26_lystate | 0.0901 | 39.4851 | 6.2837 | 5.2636 | 0.0798 | 35.8998 | 5.9916 | 4.8417 |
| HAOEC | 0.6683 | 18.0296 | 4.2461 | 3.3848 | 0.6677 | 17.5784 | 4.1927 | 3.1416 |
| HEK293T | 0.3765 | 16.5299 | 4.0657 | 3.1876 | 0.6461 | 11.0681 | 3.3269 | 2.6517 |
| HL60 | 0.4633 | 11.4203 | 3.3794 | 2.7111 | 0.6226 | 9.6282 | 3.1029 | 2.5825 |
| HaCaT | 0.2289 | 44.3847 | 6.6622 | 4.8588 | 0.2811 | 39.1072 | 6.2536 | 4.9183 |
| HepG2 | 0.428 | 31.5882 | 5.6203 | 4.3371 | 0.05 | 39.9588 | 6.3213 | 5.1588 |
| Jurkat | 0.4148 | 24.0002 | 4.899 | 3.8913 | 0.3752 | 24.4667 | 4.9464 | 4.2259 |
| K562 | -1 | 23.8188 | 4.8804 | 4.6064 | -1 | 8.9454 | 2.9909 | 2.624 |
| Mus_musculus_BMDC_lystate | 0.3239 | 10.8574 | 3.2951 | 2.4241 | 0.3697 | 10.7146 | 3.2733 | 2.3409 |
| Mus_musculus_liver_lystate | -0.1671 | 36.438 | 6.0364 | 4.8889 | 0.0819 | 32.2965 | 5.683 | 4.659 |
| Oleispira_antarctica_RB-8_lystate_R1 | 0.0454 | 26.8454 | 5.1813 | 3.8702 | 0.0335 | 18.6379 | 4.3172 | 3.2358 |
| Picrophilus_torridus_DSM9790_lystate | 0.165 | 41.0108 | 6.404 | 4.9941 | 0.2599 | 24.9528 | 4.9953 | 3.9412 |
| Saccharomyces_cerevisiae_lystate | 0.2978 | 17.0026 | 4.1234 | 3.1669 | 0.3283 | 17.5996 | 4.1952 | 3.1873 |
| Thermus_thermophilus_HB27_cells | 0.0117 | 1354.8 | 36.8076 | 29.4928 | 0.1002 | 20153.3 | 141.962 | 25.2994 |
| Thermus_thermophilus_HB27_lystate | 0.1008 | 59.3835 | 7.7061 | 6.2203 | 0.2217 | 36.9322 | 6.0772 | 4.8796 |
| U937 | 0.2618 | 18.036 | 4.2469 | 3.2158 | 0.6588 | 11.882 | 3.447 | 2.6935 |
| colon_cancer_spheroids | -0.0616 | 21.4252 | 4.6287 | 3.6292 | 0.1363 | 17.5956 | 4.1947 | 3.4264 |
| pTcells | 1 | 27.8208 | 5.2745 | 5.2123 | -0.5 | 8.1613 | 2.8568 | 2.5818 |
| MEANS | 0.230976 | 99.4846 | 7.28344 | 5.42048 | 0.231708 | 835.74 | 10.5108 | 4.86684 |

Table 9: Assessment of methodologies employing sequence-based embeddings from ESM2: The performance of individual models was assessed for each species and subsequently tested on that target species inside the FLIP partition.
| Methods | MSE |  |  |  | rankN |  |  |  |
| --- | --- | --- | --- | --- | --- | --- | --- | --- |
|  | Spearman | MSE | RMSE | MAE | Spearman | MSE | RMSE | MAE |
| Arabidopsis_thaliana_seedling_lystate | 0.2757 | 31.0004 | 5.5678 | 4.3043 | 0.1801 | 33.4706 | 5.7854 | 4.4929 |
| Bacillus_subtilis_168_lystate_R1 | 0.2119 | 26.0025 | 5.0993 | 3.944 | -0.0782 | 26.5269 | 5.1504 | 3.9686 |
| Caenorhabditis_elegans_lystate | 0.4443 | 251.104 | 15.8463 | 4.3104 | 0.4126 | 23.4481 | 4.8423 | 3.6256 |
| Danio_rerio_Zenodo_lystate | 0.1636 | 113.312 | 10.6448 | 8.979 | 0.4818 | 369.521 | 19.2229 | 18.4969 |
| Drosophila_melanogaster_SII_lystate | 0.3183 | 11.3899 | 3.3749 | 2.6573 | 0.1157 | 11.7702 | 3.4308 | 2.7413 |
| Ecoli_Lysate | 0.4357 | 35.479 | 5.9564 | 4.8229 | 0.4253 | 42.1144 | 6.4896 | 5.3407 |
| Escherichia_coli_cells | 0.5606 | 32.7802 | 5.7254 | 4.8546 | 0.3895 | 70.6396 | 8.4047 | 7.2346 |
| Geobacillus_stearothermophilus_NCA26_lystate | 0.2564 | 31.6178 | 5.623 | 4.5296 | 0.1179 | 34.9162 | 5.909 | 4.8523 |
| HAOEC | 0.6832 | 16.7097 | 4.0877 | 3.2149 | 0.6621 | 20.0385 | 4.4764 | 3.2012 |
| HEK293T | 0.5759 | 12.533 | 3.5402 | 2.7081 | 0.6298 | 12.7439 | 3.5699 | 2.8206 |
| HL60 | 0.6247 | 8.6127 | 2.9347 | 2.4174 | 0.6328 | 12.4881 | 3.5338 | 2.8032 |
| HaCaT | 0.6781 | 23.5524 | 4.8531 | 3.6007 | 0.2961 | 81.834 | 9.0462 | 7.3093 |
| HepG2 | 0.4166 | 27.5064 | 5.2447 | 4.198 | 0.7306 | 18.2769 | 4.2751 | 3.3763 |
| Jurkat | 0.5497 | 20.3893 | 4.5155 | 3.6091 | 0.56 | 26.8047 | 5.1773 | 4.1779 |
| K562 | -1 | 20.0878 | 4.4819 | 3.5169 | -1 | 33.2189 | 5.7636 | 5.5281 |
| Mus_musculus_BMDC_lystate | 0.3265 | 10.8 | 3.2863 | 2.3875 | 0.0678 | 12.1015 | 3.4787 | 2.5353 |
| Mus_musculus_liver_lystate | 0.094 | 31.4523 | 5.6082 | 4.6244 | 0.0873 | 29.042 | 5.3891 | 4.4974 |
| Oleispira_antarctica_RB-8_lystate_R1 | 0.0911 | 18.3544 | 4.2842 | 3.2036 | -0.0215 | 109.501 | 10.4643 | 8.8855 |
| Picrophilus_torridus_DSM9790_lystate | 0.222 | 29.4879 | 5.4303 | 4.3721 | 0.0355 | 234.615 | 15.3172 | 13.448 |
| Saccharomyces_cerevisiae_lystate | 0.2468 | 18.84 | 4.3405 | 3.2997 | 0.1777 | 17.2934 | 4.1585 | 3.3133 |
| Thermus_thermophilus_HB27_cells | 0.1717 | 57.7222 | 7.5975 | 6.3657 | 0.1418 | 50.6287 | 7.1154 | 5.9142 |
| Thermus_thermophilus_HB27_lystate | 0.0034 | 50.3409 | 7.0951 | 5.8207 | 0.1577 | 38.7479 | 6.2248 | 5.1955 |
| U937 | 0.0412 | 51.165 | 7.153 | 4.7931 | 0.5824 | 23.3845 | 4.8357 | 3.8638 |
| colon_cancer_spheroids | 0.1359 | 19.8405 | 4.4543 | 3.6356 | -0.02 | 22.0879 | 4.6998 | 3.6528 |
| pTcells | 0.5 | 3.8066 | 1.9511 | 1.9092 | 1 | 193.474 | 13.9095 | 13.5656 |
| MEANS | 0.281092 | 38.1555 | 5.54785 | 4.08315 | 0.270592 | 61.9475 | 6.82682 | 5.79364 |

**Table 10:** Assessment of methodologies employing embeddings from protein structures using Inverse Folding algorithms (PiFold): The performance of individual models was assessed for each species and subsequently tested on that target species inside the FLIP partition. Spearman correlation, mean square error (MSE), root mean square error (RMSE) and mean absolute error (MAE). These metrics were used to analyse the effectiveness of the four methods investigated in this study: balanceMSE, biasg, MSE and RankN.

| Methods | balanceMSE |  |  |  | biasg |  |  |  |
| --- | --- | --- | --- | --- | --- | --- | --- | --- |
|  | Spearman | MSE | RMSE | MAE | Spearman | MSE | RMSE | MAE |
| Arabidopsis_thaliana_seedling_lystate | 0.1901 | 33.1926 | 5.7613 | 4.416 | 0.2502 | 31.8047 | 5.6396 | 4.36 |
| Bacillus_subtilis_168_lystate_R1 | 0.235 | 26.0891 | 5.1078 | 3.8833 | 0.2005 | 26.9053 | 5.187 | 4.0578 |
| Caenorhabditis_elegans_lystate | 0.2814 | 23.3374 | 4.8309 | 3.8388 | 0.2717 | 24.0761 | 4.9067 | 3.7928 |
| Danio_rerio_Zenodo_lystate | 0.0545 | 55.0791 | 7.4215 | 5.9855 | -0.1909 | 85.0409 | 9.2218 | 6.6883 |
| Drosophila_melanogaster_SII_lystate | 0.332 | 11.2 | 3.3466 | 2.6635 | 0.2971 | 11.8738 | 3.4458 | 2.6509 |
| Ecoli_Lysate | 0.2438 | 49.9904 | 7.0704 | 5.551 | 0.3893 | 38.8821 | 6.2355 | 5.0328 |
| Escherichia_coli_cells | 0.2479 | 62.1012 | 7.8804 | 6.5516 | 0.1184 | 53.963 | 7.346 | 6.2021 |
| Geobacillus_stearothermophilus_NCA26_lystate | 0.0345 | 43.1849 | 6.5715 | 5.3951 | 0.0842 | 41.817 | 6.4666 | 5.1216 |
| HAOEC | 0.5553 | 21.1289 | 4.5966 | 3.6297 | 0.5702 | 21.5063 | 4.6375 | 3.5582 |
| HEK293T | 0.3274 | 18.916 | 4.3493 | 3.4813 | 0.3553 | 16.8959 | 4.1105 | 3.3724 |
| HL60 | 0.5285 | 12.7037 | 3.5642 | 2.8803 | 0.5094 | 12.7012 | 3.5639 | 2.8331 |
| HaCaT | 0.0953 | 46.8168 | 6.8423 | 5.0379 | 0.0528 | 57.0223 | 7.5513 | 5.5844 |
| HepG2 | 0.4819 | 27.8663 | 5.2789 | 4.0159 | 0.5148 | 25.9791 | 5.097 | 4.1233 |
| Jurkat | 0.3438 | 27.167 | 5.2122 | 4.2192 | 0.0998 | 29.5164 | 5.4329 | 4.4984 |
| K562 | -1 | 20.4271 | 4.5196 | 4.0903 | -1 | 14.7407 | 3.8394 | 3.6112 |
| Mus_musculus_BMDC_lystate | 0.1828 | 12.4816 | 3.5329 | 2.5956 | 0.2007 | 12.7979 | 3.5774 | 2.5749 |
| Mus_musculus_liver_lystate | 0.1398 | 29.6782 | 5.4478 | 4.4864 | 0.1861 | 139.129 | 11.7953 | 5.8553 |
| Oleispira_antartica_RB-8_lystate_R1 | -0.0207 | 24.0207 | 4.9011 | 3.762 | 0.0777 | 17.7828 | 4.217 | 3.183 |
| Picrophilus_torridus_DSM9790_lystate | 0.0646 | 37.0564 | 6.0874 | 4.8446 | 0.113 | 28.2721 | 5.3171 | 4.3029 |
| Saccharomyces_cerevisiae_lystate | 0.0481 | 19.6548 | 4.4334 | 3.4708 | 0.1198 | 18.515 | 4.3029 | 3.395 |
| Thermus_thermophilus_HB27_cells | -0.0552 | 73.09 | 8.5493 | 7.1818 | -0.0566 | 63.1341 | 7.9457 | 6.5063 |
| Thermus_thermophilus_HB27_lystate | 0.0847 | 45.8389 | 6.7704 | 5.6529 | 0.1028 | 41.072 | 6.4087 | 5.3268 |
| U937 | 0.5971 | 15.4958 | 3.9365 | 2.7684 | 0.1588 | 17.6512 | 4.2013 | 3.4349 |
| colon_cancer_spheroids | 0.1604 | 17.373 | 4.1681 | 3.4119 | 0.0388 | 19.2596 | 4.3886 | 3.5495 |
| pTcells | 0.5 | 26.5078 | 5.1486 | 4.3754 | 0.5 | 8.3234 | 2.885 | 2.6848 |
| MEANS | 0.18612 | 31.2159 | 5.41316 | 4.32757 | 0.158556 | 34.3465 | 5.50882 | 4.25203 |

Table 10: Assessment of methodologies employing embeddings from protein structures using Inverse Folding algorithms (PiFold): The performance of individual models was assessed for each species and subsequently tested on that target species inside the FLIP partition.
| Methods | MSE |  |  |  | rankN |  |  |  |
| --- | --- | --- | --- | --- | --- | --- | --- | --- |
|  | Spearman | MSE | RMSE | MAE | Spearman | MSE | RMSE | MAE |
| Arabidopsis_thaliana_seedling_lystate | 0.1513 | 33.8696 | 5.8198 | 4.4579 | 0.1197 | 32.8702 | 5.7332 | 4.5013 |
| Bacillus_subtilis_168_lystate_R1 | 0.1527 | 28.1217 | 5.303 | 4.1673 | 0.147 | 26.1166 | 5.1104 | 4.0038 |
| Caenorhabditis_elegans_lystate | 0.3198 | 24.1727 | 4.9166 | 3.7621 | 0.1201 | 25.0226 | 5.0023 | 3.9697 |
| Danio_rerio_Zenodo_lystate | 0.6 | 33.9685 | 5.8282 | 4.6552 | 0.1545 | 39.9804 | 6.323 | 4.7511 |
| Drosophila_melanogaster_SII_lystate | 0.3204 | 10.97 | 3.3121 | 2.5842 | 0.3396 | 12.9318 | 3.5961 | 2.8986 |
| Ecoli_Lysate | 0.2583 | 42.3656 | 6.5089 | 5.2799 | 0.4019 | 48.3124 | 6.9507 | 5.6156 |
| Escherichia_coli_cells | 0.367 | 58.9055 | 7.675 | 6.5599 | -0.0351 | 61.2983 | 7.8293 | 6.7248 |
| Geobacillus_stearothermophilus_NCA26_lystate | -0.106 | 39.5683 | 6.2903 | 4.9254 | 0.1681 | 33.9255 | 5.8246 | 4.5943 |
| HAOEC | 0.6131 | 19.766 | 4.4459 | 3.5534 | 0.5151 | 24.2065 | 4.92 | 3.6582 |
| HEK293T | 0.5067 | 14.7227 | 3.837 | 3.0679 | 0.4558 | 14.4314 | 3.7989 | 2.998 |
| HL60 | 0.5877 | 9.8337 | 3.1359 | 2.5131 | 0.1545 | 16.2364 | 4.0294 | 3.0754 |
| HaCaT | 0.4174 | 34.1148 | 5.8408 | 4.4338 | 0.3616 | 36.3923 | 6.0326 | 4.6482 |
| HepG2 | 0.6413 | 19.7246 | 4.4412 | 3.4868 | 0.5916 | 23.1588 | 4.8124 | 3.8102 |
| Jurkat | 0.3163 | 26.6553 | 5.1629 | 4.2983 | 0.271 | 24.4258 | 4.9422 | 4.2085 |
| K562 | 1 | 27.3812 | 5.2327 | 5.1922 | -1 | 5.3537 | 2.3138 | 1.9069 |
| Mus_musculus_BMDC_lystate | 0.2036 | 12.4449 | 3.5277 | 2.4947 | 0.1087 | 11.8251 | 3.4388 | 2.535 |
| Mus_musculus_liver_lystate | 0.2019 | 26.4774 | 5.1456 | 4.1962 | -0.0673 | 30.653 | 5.5365 | 4.6092 |
| Oleispira_antartica_RB-8_lystate_R1 | 0.0616 | 18.4959 | 4.3007 | 3.3799 | 0.1153 | 16.5094 | 4.0632 | 3.0985 |
| Picrophilus_torridus_DSM9790_lystate | 0.0001 | 31.7089 | 5.6311 | 4.547 | 0.2134 | 31.4108 | 5.6045 | 4.391 |
| Saccharomyces_cerevisiae_lystate | 0.2475 | 17.3746 | 4.1683 | 3.1798 | 0.2441 | 23.5091 | 4.8486 | 3.9765 |
| Thermus_thermophilus_HB27_cells | -0.015 | 58.7667 | 7.6659 | 6.0328 | -0.057 | 48.9728 | 6.9981 | 5.7465 |
| Thermus_thermophilus_HB27_lystate | 0.0471 | 49.1931 | 7.0138 | 5.7324 | 0.1022 | 37.4915 | 6.123 | 5.083 |
| U937 | -0.1235 | 24.0018 | 4.8992 | 3.9386 | -0.1324 | 18.1745 | 4.2632 | 3.5649 |
| colon_cancer_spheroids | 0.0761 | 18.1357 | 4.2586 | 3.4174 | 0.0622 | 17.8032 | 4.2194 | 3.4283 |
| pTcells | 0.5 | 26.305 | 5.1288 | 4.9994 | -0.5 | 18.0692 | 4.2508 | 3.7284 |
| MEANS | 0.293816 | 28.2818 | 5.1796 | 4.19422 | 0.114184 | 27.1633 | 5.0626 | 4.06104 |

**Table 11:** Assessment of methodologies using the combination of ESM2 and PiFold embeddings: The performance of individual models was assessed for each species and subsequently tested on that target species inside the FLIP partition. Spearman correlation, mean square error (MSE), root mean square error (RMSE) and mean absolute error (MAE). These metrics were used to analyse the effectiveness of the four methods investigated in this study: balanceMSE, biasg, MSE and RankN.

| Methods | balanceMSE |  |  |  | biasg |  |  |  |
| --- | --- | --- | --- | --- | --- | --- | --- | --- |
|  | Spearman | MSE | RMSE | MAE | Spearman | MSE | RMSE | MAE |
| Arabidopsis_thaliana_seedling_lystate | 0.3505 | 29.5605 | 5.437 | 4.1579 | 0.3321 | 30.1566 | 5.4915 | 4.1206 |
| Bacillus_subtilis_168_lystate_R1 | 0.2232 | 25.9268 | 5.0918 | 3.901 | 0.3015 | 26.9116 | 5.1876 | 4.2273 |
| Caenorhabditis_elegans_lystate | 0.368 | 22.4169 | 4.7346 | 3.6779 | 0.4724 | 20.5988 | 4.5386 | 3.5272 |
| Danio_rerio_Zenodo_lystate | -0.2545 | 72.4703 | 8.5129 | 6.3495 | 0.3364 | 48.9019 | 6.993 | 5.3776 |
| Drosophila_melanogaster_SII_lystate | 0.3374 | 10.9702 | 3.3121 | 2.6325 | 0.4173 | 10.1633 | 3.188 | 2.5342 |
| Ecoli_lystate | 0.382 | 40.7093 | 6.3804 | 5.0832 | 0.5134 | 31.3316 | 5.5975 | 4.5145 |
| Escherichia_coli_cells | 0.5293 | 44.9233 | 6.7025 | 5.6516 | 0.5683 | 34.6805 | 5.889 | 4.8422 |
| Geobacillus_stearothermophilus_NCA26_lystate | 0.1632 | 39.2467 | 6.2647 | 5.0332 | 0.2924 | 31.5223 | 5.6145 | 4.5322 |
| HAOEC | 0.6825 | 16.6991 | 4.0865 | 2.9802 | 0.6785 | 16.2852 | 4.0355 | 3.166 |
| HEK293T | 0.6503 | 10.8033 | 3.2868 | 2.6429 | 0.5679 | 12.6298 | 3.5538 | 2.8093 |
| HL60 | 0.663 | 9.4643 | 3.0764 | 2.4607 | 0.663 | 9.4511 | 3.0743 | 2.5043 |
| HaCaT | -0.0057 | 47.7258 | 6.9084 | 5.1539 | -0.3245 | 48.5773 | 6.9697 | 5.5694 |
| HepG2 | 0.6987 | 16.7001 | 4.0866 | 3.242 | 0.7914 | 14.4484 | 3.8011 | 3.0977 |
| Jurkat | 0.444 | 23.4916 | 4.8468 | 3.7271 | 0.6486 | 16.1099 | 4.0137 | 3.2487 |
| K562 | -1 | 11.0728 | 3.3276 | 2.8619 | -1 | 3.0531 | 1.7473 | 1.3394 |
| Mus_musculus_BMDC_lystate | 0.3702 | 10.9503 | 3.3091 | 2.3 | 0.286 | 11.6081 | 3.4071 | 2.3945 |
| Mus_musculus_liver_lystate | 0.2139 | 26.6532 | 5.1627 | 4.2005 | -0.0108 | 31.4768 | 5.6104 | 4.6453 |
| Oleispira_antarctica_RB-8_lystate_R1 | 0.035 | 24.5496 | 4.9548 | 3.5839 | 0.1266 | 16.7649 | 4.0945 | 3.0388 |
| Picrophilus_torridus_DSM9790_lystate | 0.2688 | 29.8296 | 5.4617 | 4.3719 | 0.0523 | 27.8212 | 5.2746 | 4.1684 |
| Saccharomyces_cerevisiae_lystate | 0.3152 | 15.6218 | 3.9524 | 3.1088 | 0.2702 | 16.1805 | 4.0225 | 3.0964 |
| Thermus_thermophilus_HB27_cells | 0.1111 | 74.9102 | 8.6551 | 7.2418 | 0.0171 | 55.0932 | 7.4225 | 5.989 |
| Thermus_thermophilus_HB27_lystate | 0.0642 | 45.6323 | 6.7552 | 5.4227 | 0.3609 | 38.8648 | 6.2342 | 5.0251 |
| U937 | 0.6794 | 12.4133 | 3.5232 | 2.9262 | 0.6765 | 9.8029 | 3.131 | 2.6008 |
| colon_cancer_spheroids | 0.1879 | 16.9934 | 4.1223 | 3.4596 | -0.1437 | 18.6022 | 4.313 | 3.4689 |
| pTcells | 0.5 | 15.8435 | 3.9804 | 2.8804 | -0.5 | 14.422 | 3.7976 | 3.5517 |
| MEANS | 0.279104 | 27.8231 | 5.03728 | 3.96205 | 0.255752 | 23.8183 | 4.6801 | 3.73558 |

Table 11: Assessment of methodologies using the combination of ESM2 and PiFold embeddings: The performance of individual models was assessed for each species and subsequently tested on that target species inside the FLIP partition.
| Methods | MSE |  |  |  | rankN |  |  |  |
| --- | --- | --- | --- | --- | --- | --- | --- | --- |
|  | Spearman | MSE | RMSE | MAE | Spearman | MSE | RMSE | MAE |
| Arabidopsis_thaliana_seedling_lystate | 0.3461 | 29.1147 | 5.3958 | 4.1742 | 0.1444 | 33.0563 | 5.7495 | 4.4886 |
| Bacillus_subtilis_168_lystate_R1 | 0.1653 | 25.9705 | 5.0961 | 4.0067 | 0.1319 | 25.9661 | 5.0957 | 3.9435 |
| Caenorhabditis_elegans_lystate | 0.499 | 20.6519 | 4.5444 | 3.4613 | 0.4784 | 21.4948 | 4.6362 | 3.4993 |
| Danio_rerio_Zenodo_lystate | 0.0455 | 109.125 | 10.4463 | 8.6692 | 0.0364 | 193.913 | 13.9253 | 12.4161 |
| Drosophila_melanogaster_SII_lystate | 0.3202 | 12.096 | 3.4779 | 2.7011 | 0.3287 | 10.5318 | 3.2453 | 2.5795 |
| Ecoli_lystate | 0.4779 | 31.9304 | 5.6507 | 4.5741 | 0.5269 | 33.6137 | 5.7977 | 4.6787 |
| Escherichia_coli_cells | 0.5883 | 31.4554 | 5.6085 | 4.7336 | 0.5132 | 65.2721 | 8.0791 | 6.593 |
| Geobacillus_stearothermophilus_NCA26_lystate | 0.1593 | 37.1683 | 6.0966 | 5.0318 | 0.0914 | 33.9548 | 5.8271 | 4.8257 |
| HAOEC | 0.6695 | 16.8059 | 4.0995 | 3.148 | 0.6921 | 18.7967 | 4.3355 | 3.1634 |
| HEK293T | 0.4082 | 15.5122 | 3.9386 | 3.1006 | 0.5787 | 12.4972 | 3.5351 | 2.6919 |
| HL60 | 0.6704 | 7.8528 | 2.8023 | 2.3577 | 0.6633 | 12.8484 | 3.5845 | 2.8722 |
| HaCaT | 0.4248 | 26.0739 | 5.1063 | 4.0582 | -0.1043 | 150.318 | 12.2604 | 9.0783 |
| HepG2 | 0.6422 | 20.104 | 4.4837 | 3.5313 | 0.7702 | 18.014 | 4.2443 | 3.3321 |
| Jurkat | 0.558 | 18.3289 | 4.2812 | 3.419 | 0.6849 | 23.2359 | 4.8204 | 3.8469 |
| K562 | 1 | 42.885 | 6.5487 | 6.3414 | -1 | 25.5311 | 5.0528 | 4.0778 |
| Mus_musculus_BMDC_lystate | 0.3303 | 11.1626 | 3.3411 | 2.47 | 0.1396 | 12.6878 | 3.562 | 2.5788 |
| Mus_musculus_liver_lystate | 0.1039 | 28.9951 | 5.3847 | 4.4282 | 0.2819 | 30.818 | 5.5514 | 4.3435 |
| Oleispira_antarctica_RB-8_lystate_R1 | 0.0069 | 19.2697 | 4.3897 | 3.3043 | 0.2918 | 15.99 | 3.9988 | 2.9648 |
| Picrophilus_torridus_DSM9790_lystate | 0.2891 | 26.4809 | 5.146 | 4.1468 | 0.1515 | 137.6 | 11.7303 | 10.1876 |
| Saccharomyces_cerevisiae_lystate | 0.3907 | 15.1299 | 3.8897 | 2.9698 | 0.2702 | 16.3817 | 4.0474 | 3.1665 |
| Thermus_thermophilus_HB27_cells | 0.1968 | 58.7222 | 7.663 | 6.4591 | 0.1177 | 47.7578 | 6.9107 | 5.6658 |
| Thermus_thermophilus_HB27_lystate | -0.1086 | 45.6133 | 6.7538 | 5.5957 | 0.0697 | 35.6751 | 5.9729 | 4.9608 |
| U937 | 0.5059 | 12.1784 | 3.4898 | 3.0624 | 0.1618 | 29.4041 | 5.4226 | 4.2814 |
| colon_cancer_spheroids | 0.0397 | 18.7022 | 4.3246 | 3.4427 | 0.189 | 16.633 | 4.0784 | 3.201 |
| pTcells | -0.5 | 9.1506 | 3.025 | 2.8475 | 0.5 | 198.42 | 14.0862 | 13.8163 |
| MEANS | 0.329176 | 27.6192 | 4.99936 | 4.08139 | 0.268376 | 48.8165 | 6.22198 | 5.09014 |

### E Cross-species Prediction Scatterplots - Global Model Applied to Individual Species

**Figure 8:**
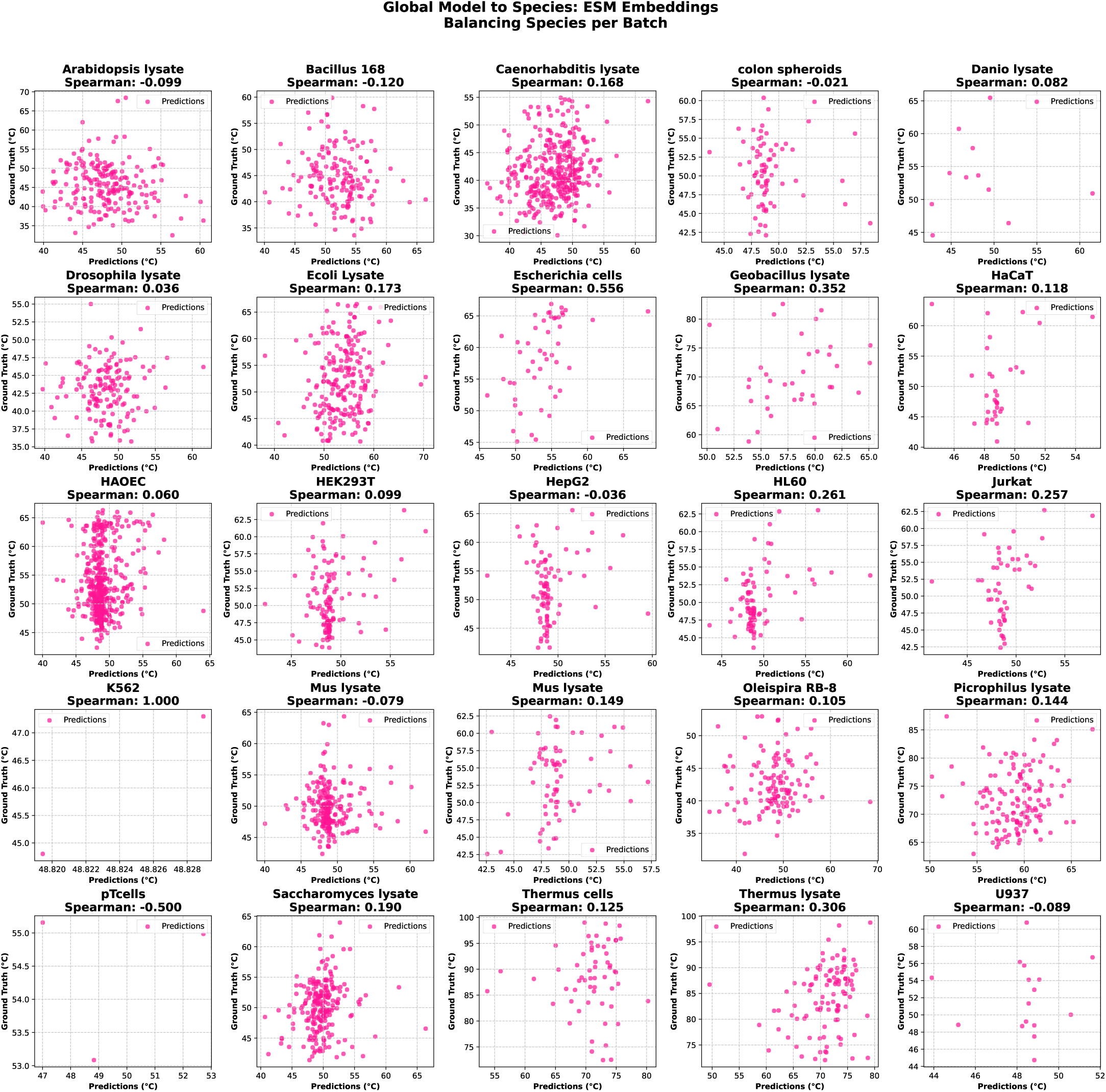
Scatterplot of global model applied to each species using balancing strategy per batch and ESM embeddings.

**Figure 9:**
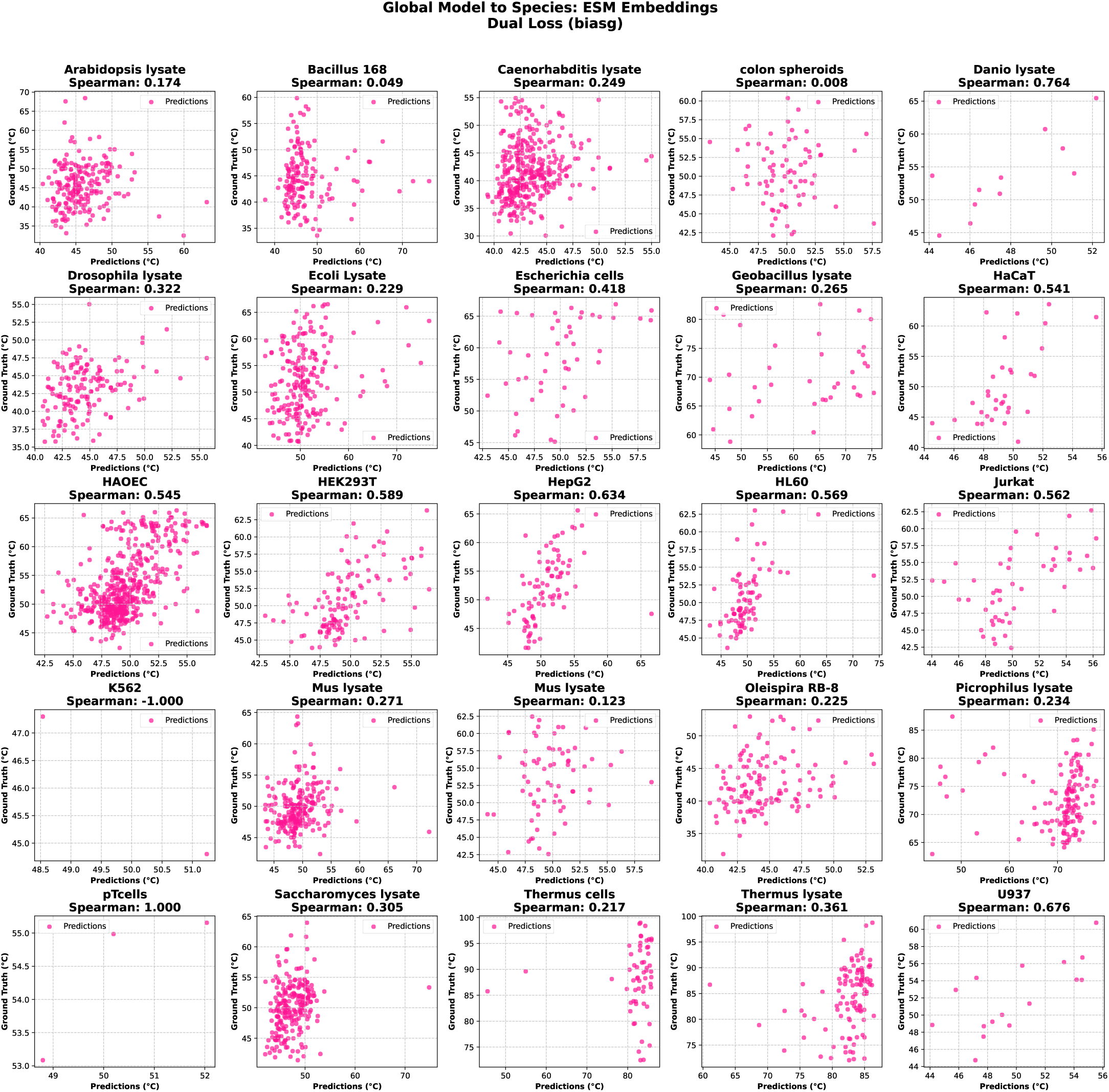
Scatterplot of global model applied to each species using dual loss function and ESM embeddings.

**Figure 10:**
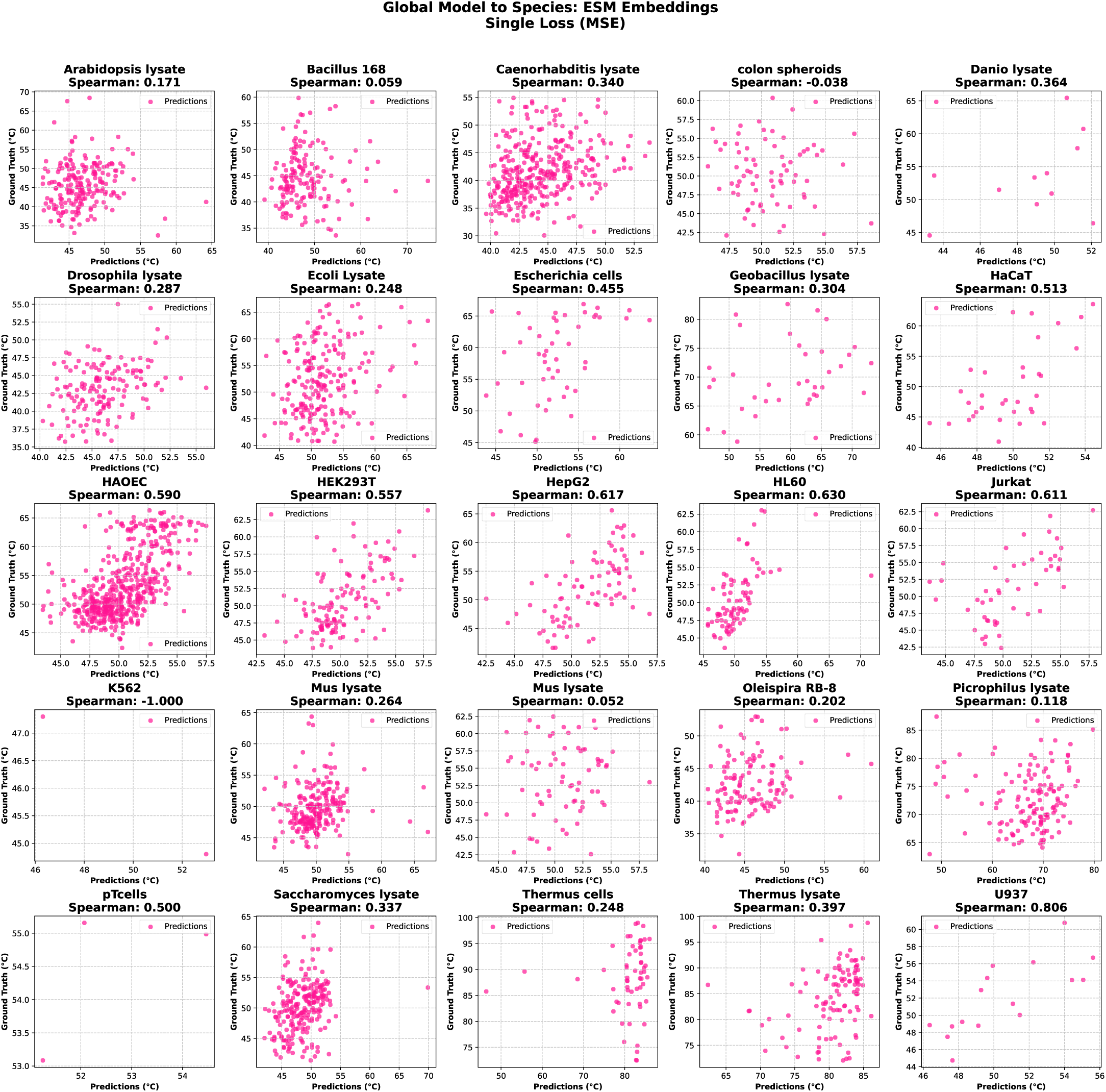
Scatterplot of global model applied to each species using MSE as loss function and ESM embeddings.

**Figure 11:**
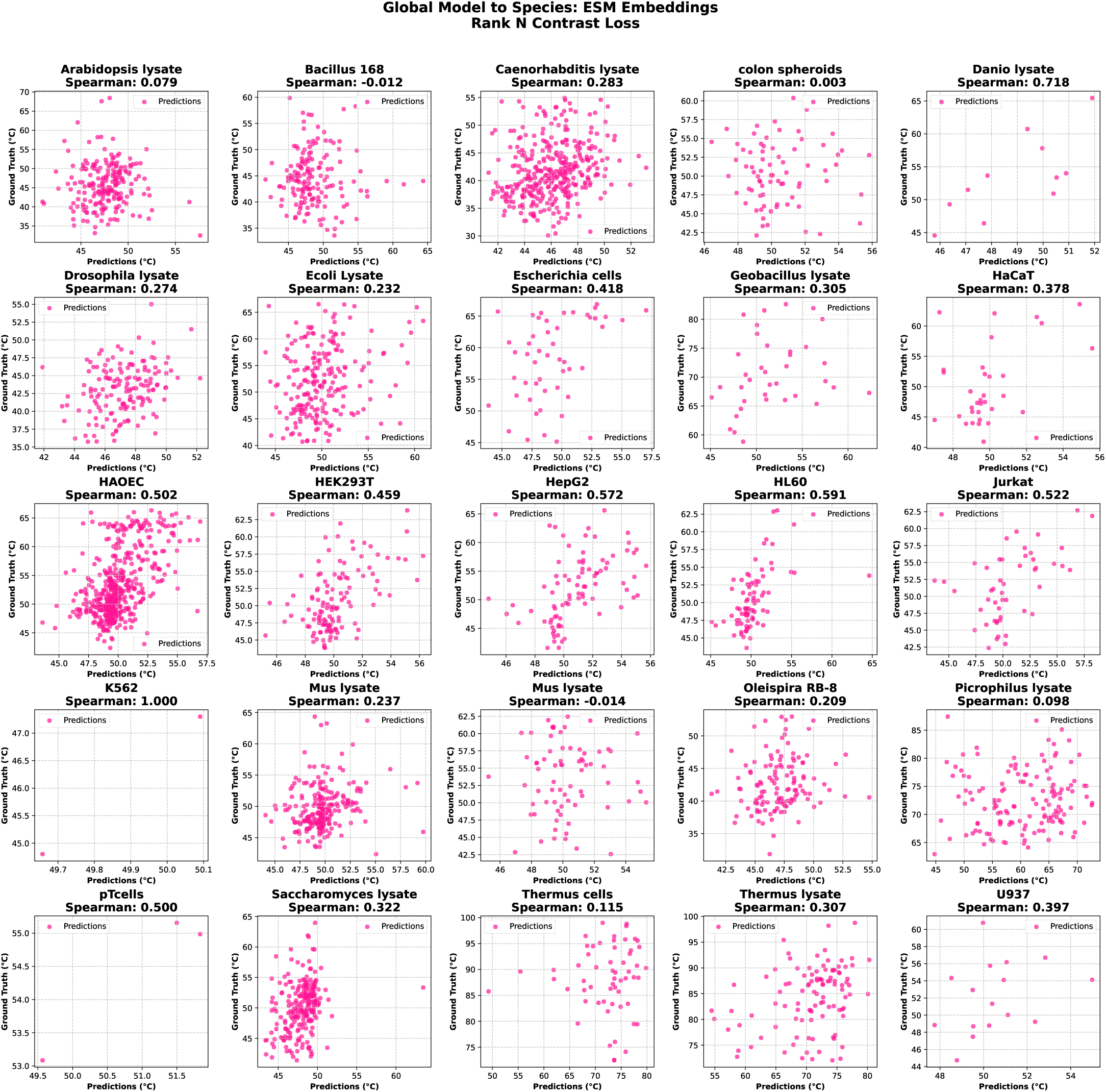
Scatterplot of global model applied to each species using Rank N Contrast loss and ESM embeddings.

**Figure 12:**
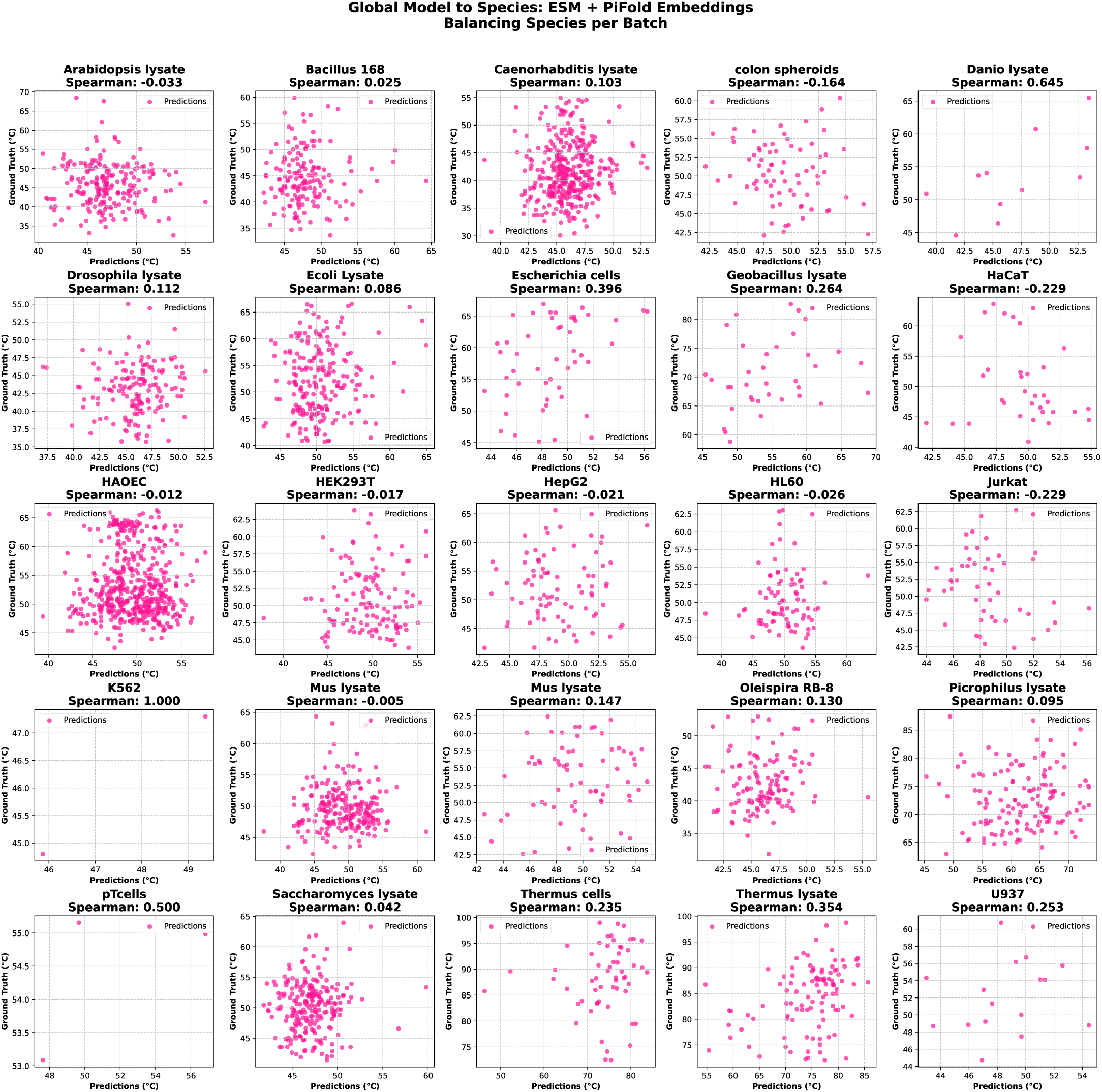
Scatterplot of global model applied to each species using balancing strategy per batch in combination with ESM and PiFold embeddings.

**Figure 13:**
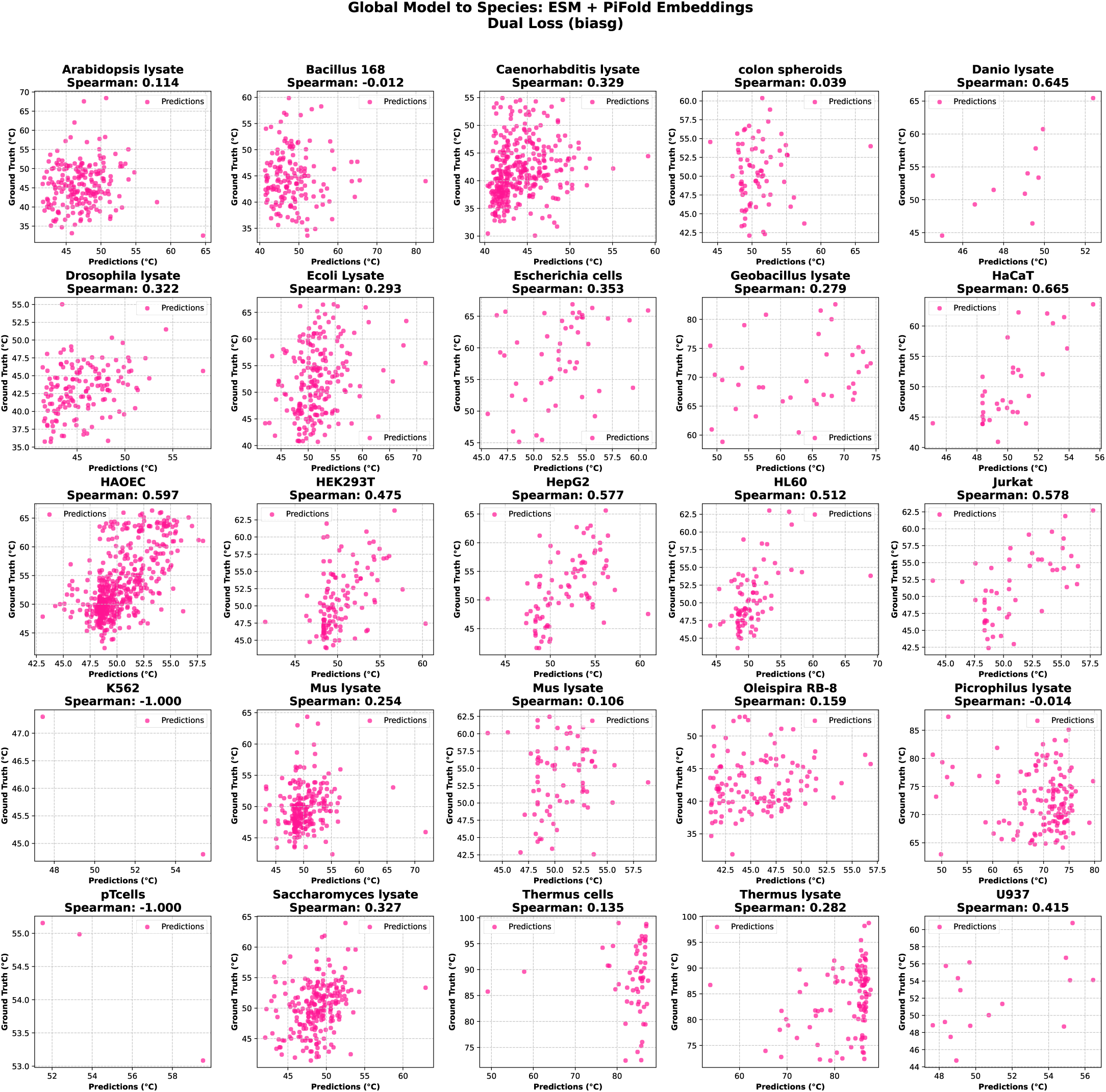
Scatterplot of global model applied to each species using dual loss function in combination with ESM and PiFold embeddings.

**Figure 14:**
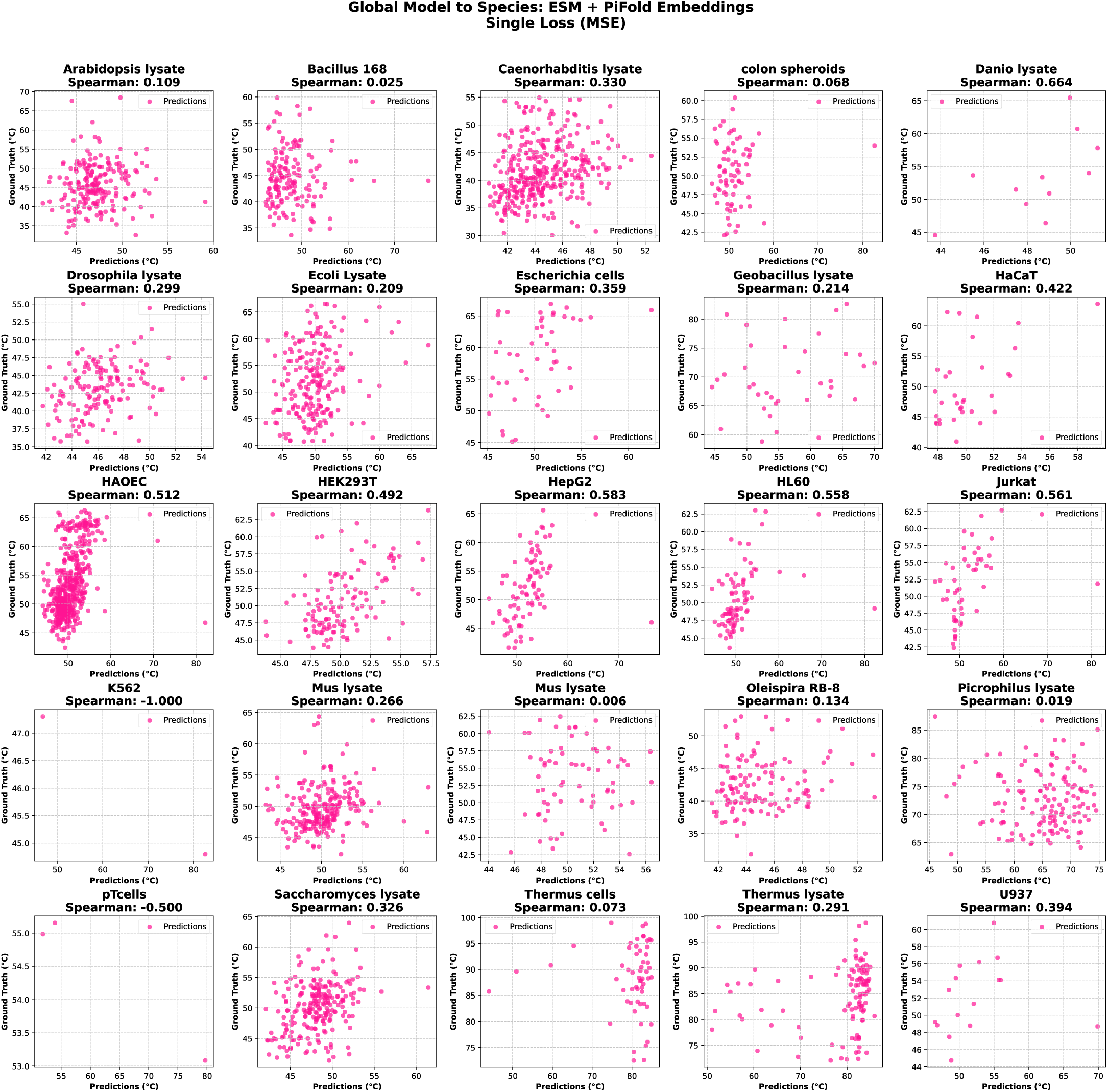
Scatterplot of global model applied to each species using MSE as loss function in combination with ESM and PiFold embeddings.

**Figure 15:**
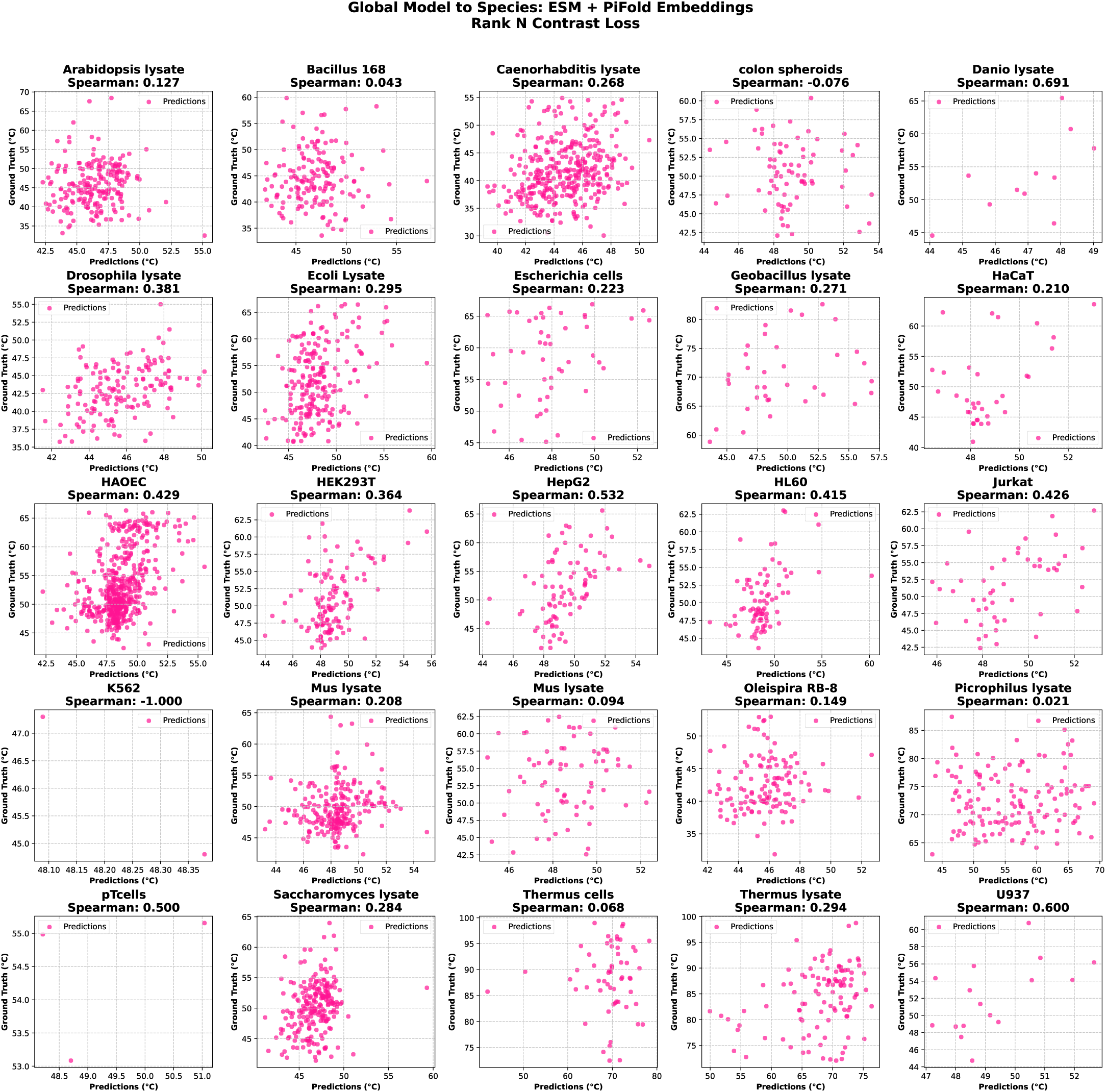
Scatterplot of global model applied to each species using Rank N Contrast loss in combination with ESM and PiFold embeddings.

**Figure 16:**
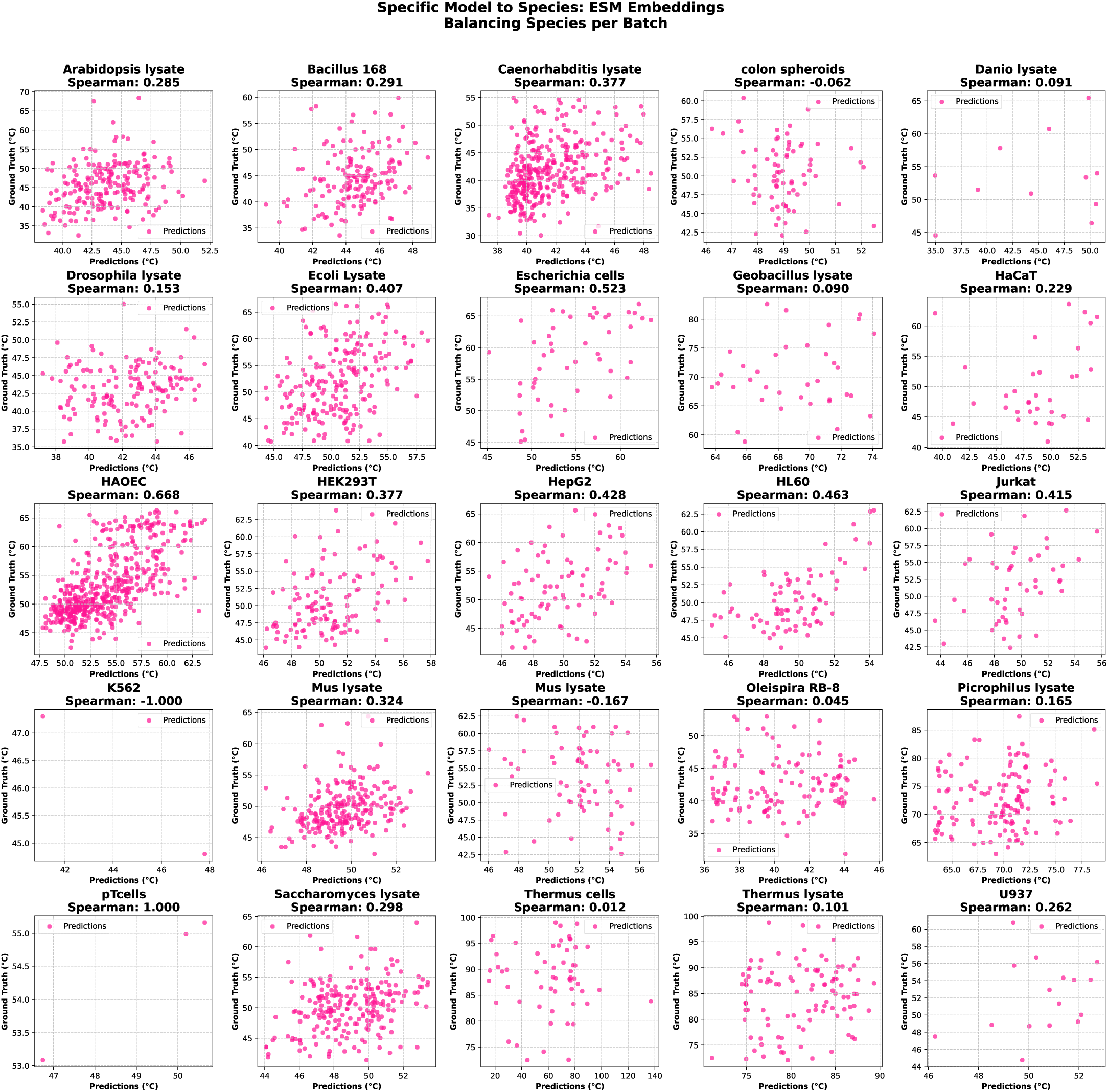
Scatterplot of individual models applied to their target species using balancing strategy per batch with ESM embeddings.

**Figure 17:**
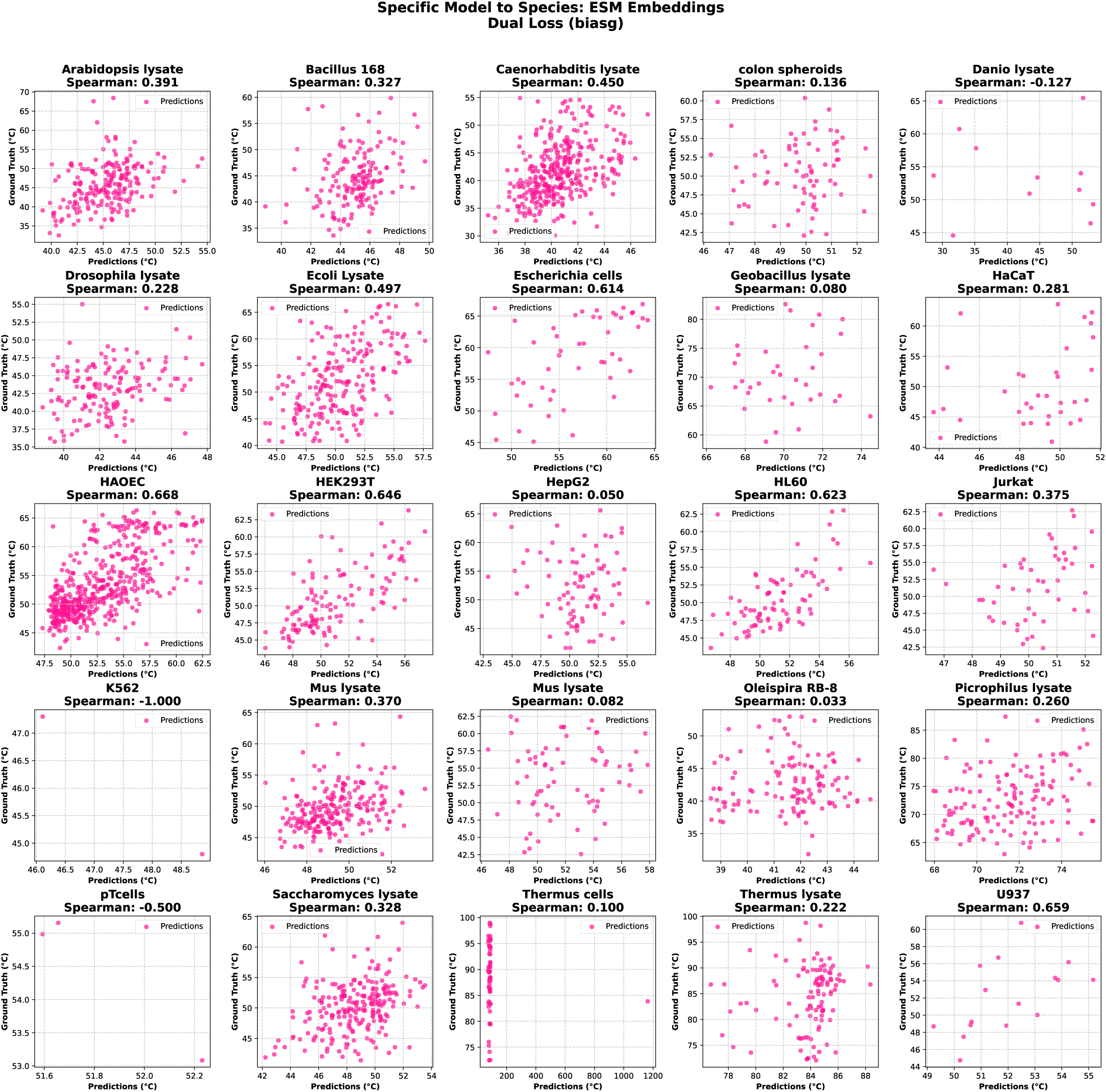
Scatterplot of individual models applied to their target species using dual loss function with ESM embeddings.

**Figure 18:**
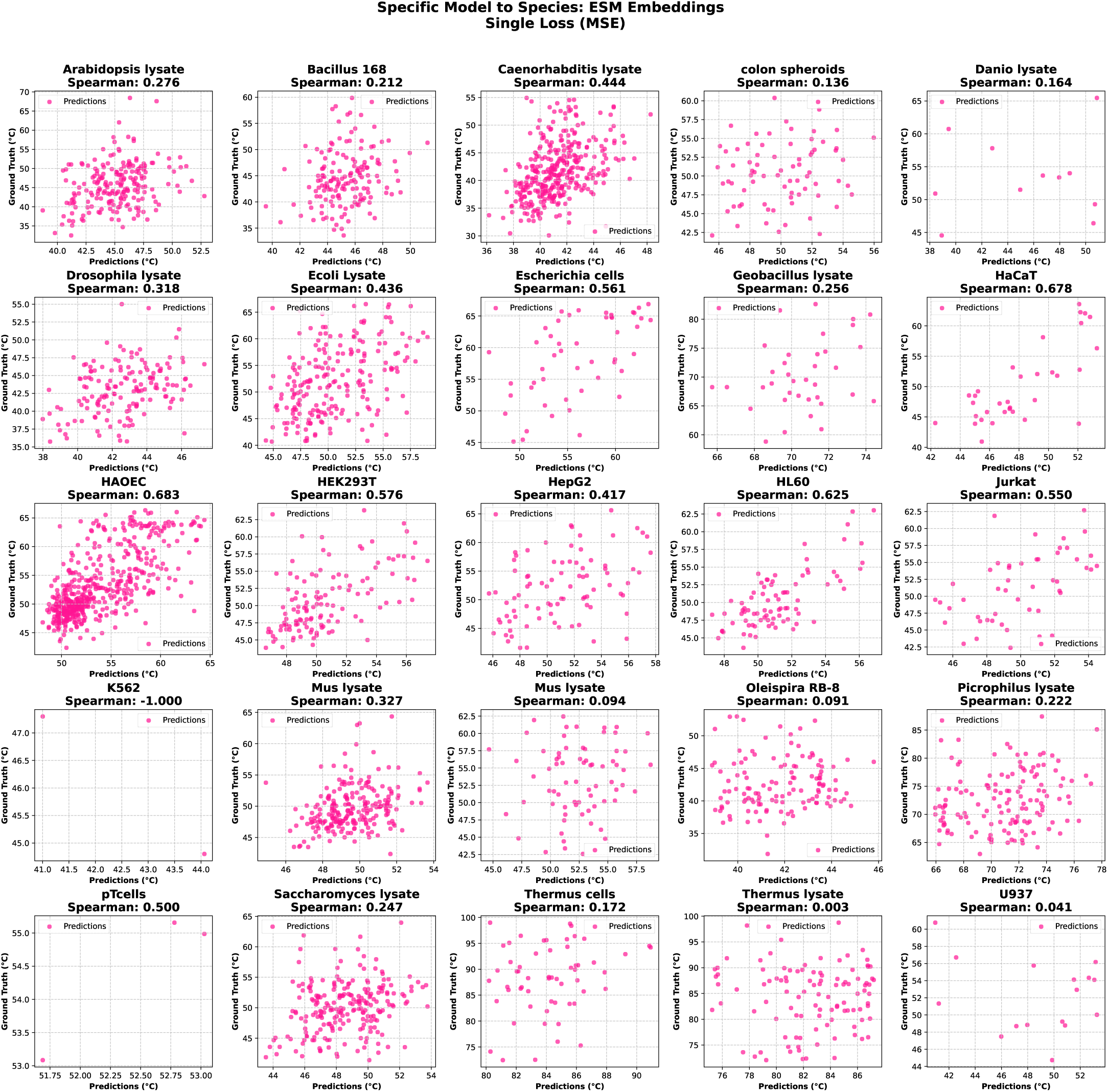
Scatterplot of individual models applied to their target species using MSE as loss function with ESM embeddings.

**Figure 19:**
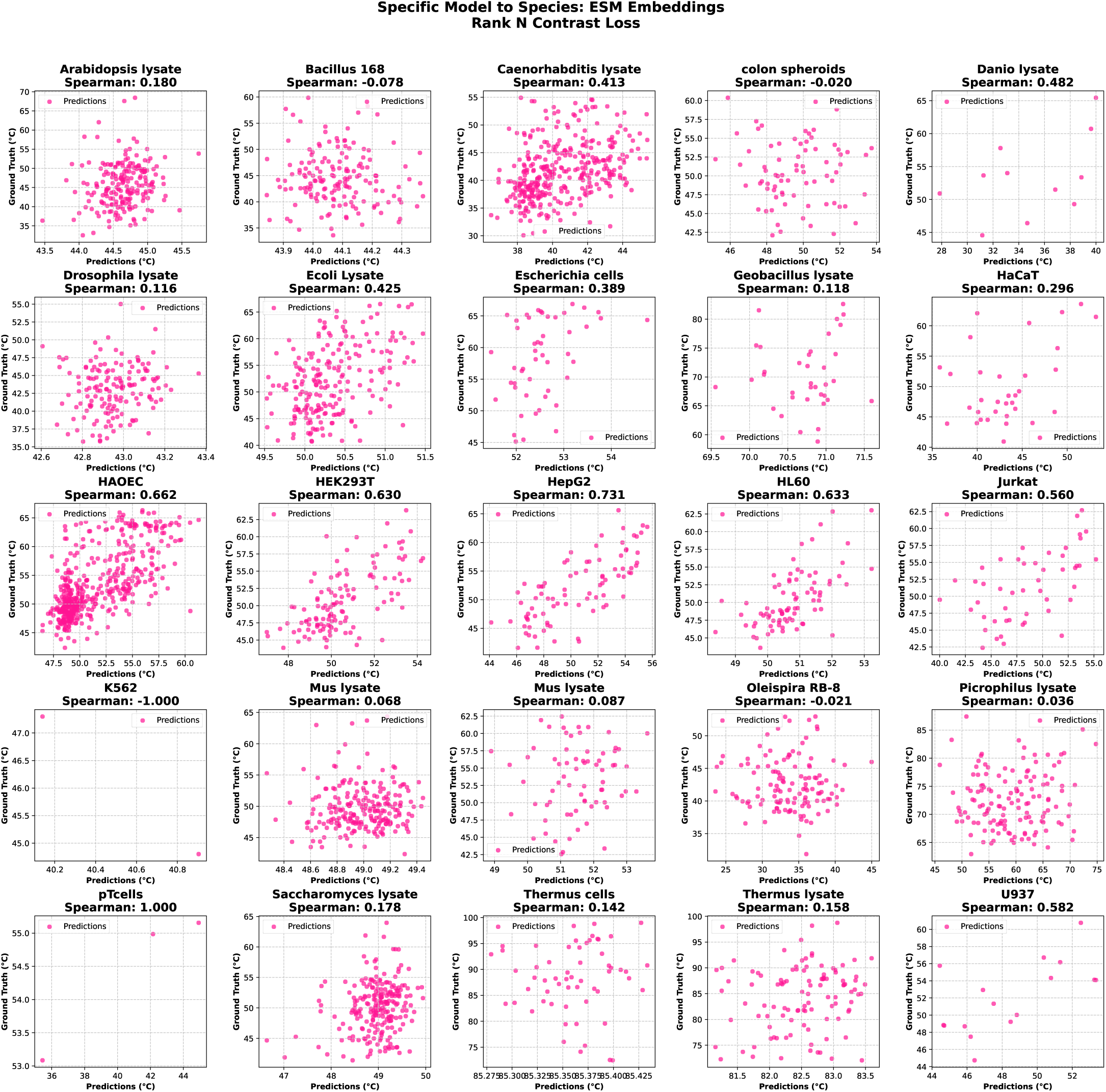
Scatterplot of individual models applied to their target species using Rank N Contrast loss with ESM embeddings.

**Figure 20:**
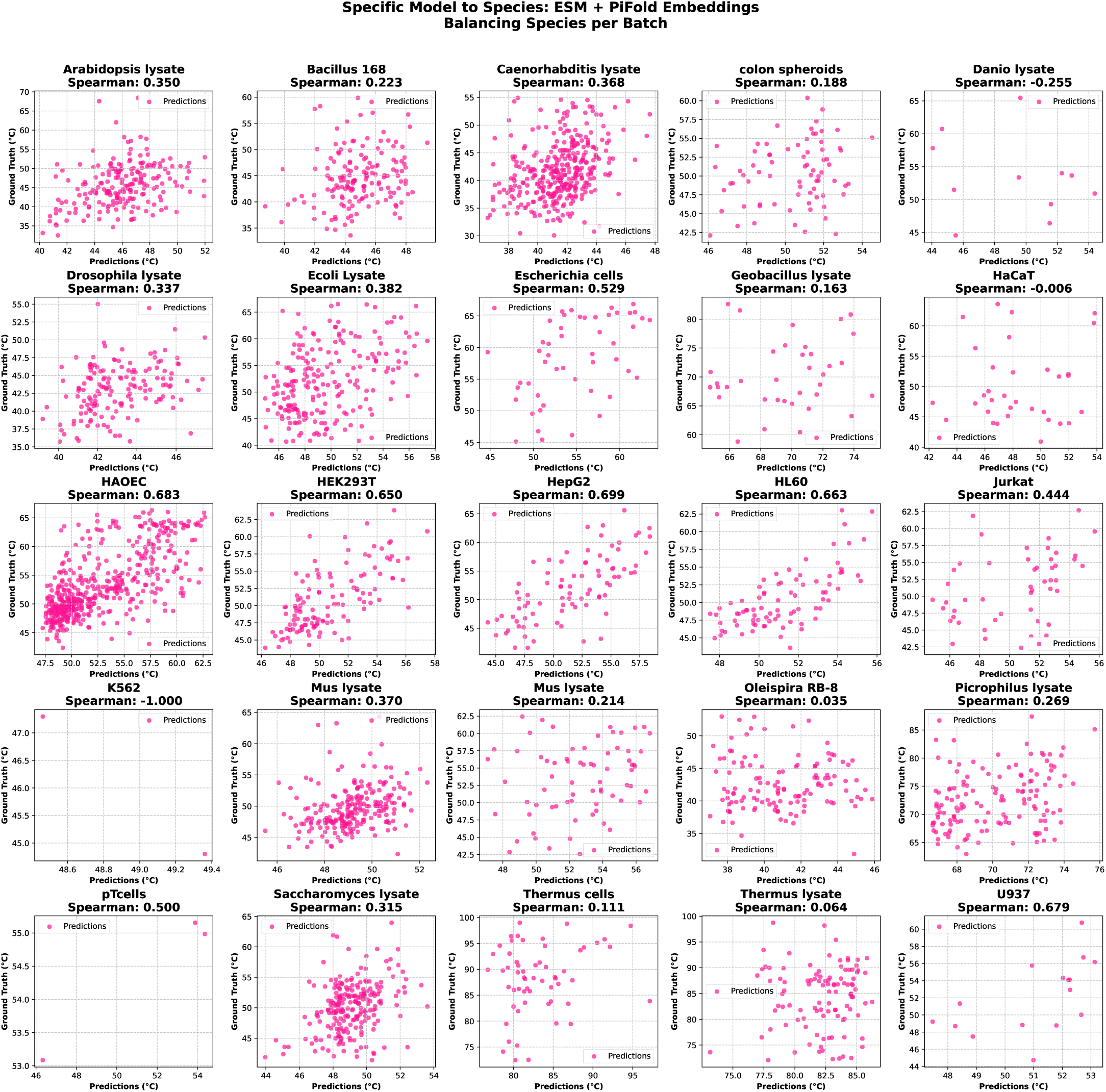
Scatterplot of individual models applied to their target species using balancing strategy per batch in combination with ESM and PiFold embeddings.

**Figure 21:**
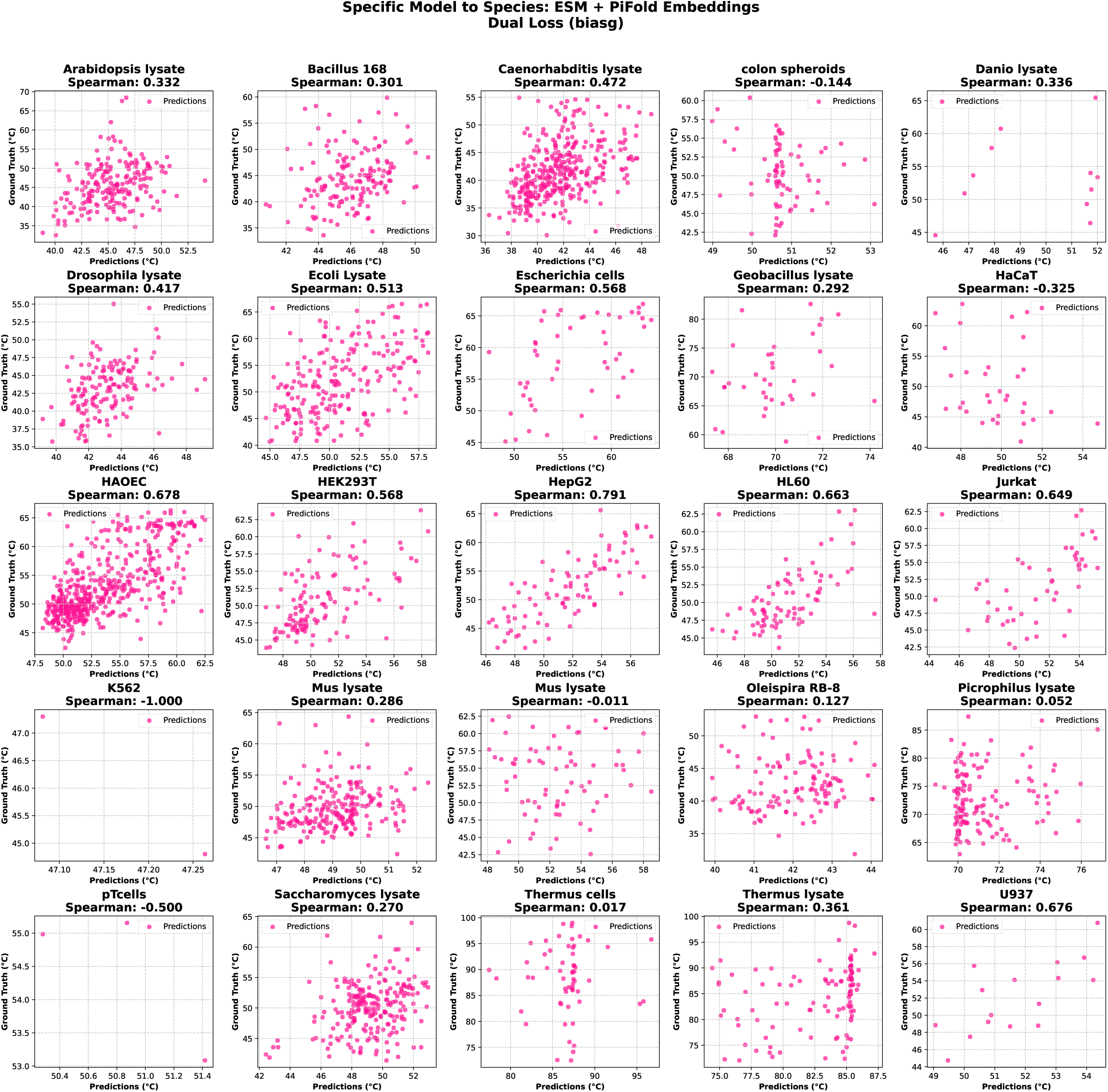
Scatterplot of individual models applied to their target species using dual loss function in combination with ESM and PiFold embeddings.

**Figure 22:**
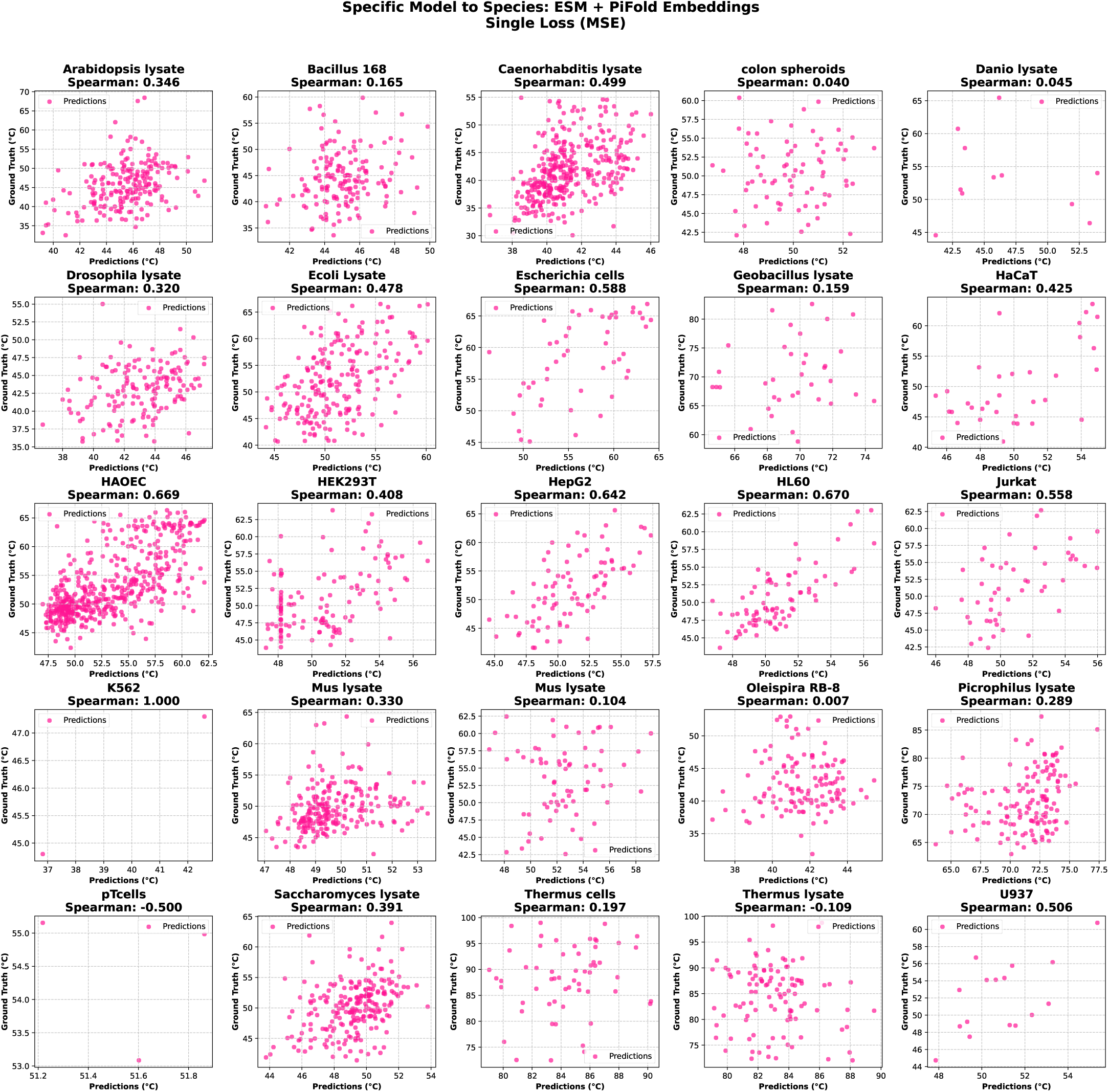
Scatterplot of individual models applied to their target species using MSE as loss function in combination with ESM and PiFold embeddings.

**Figure 23:**
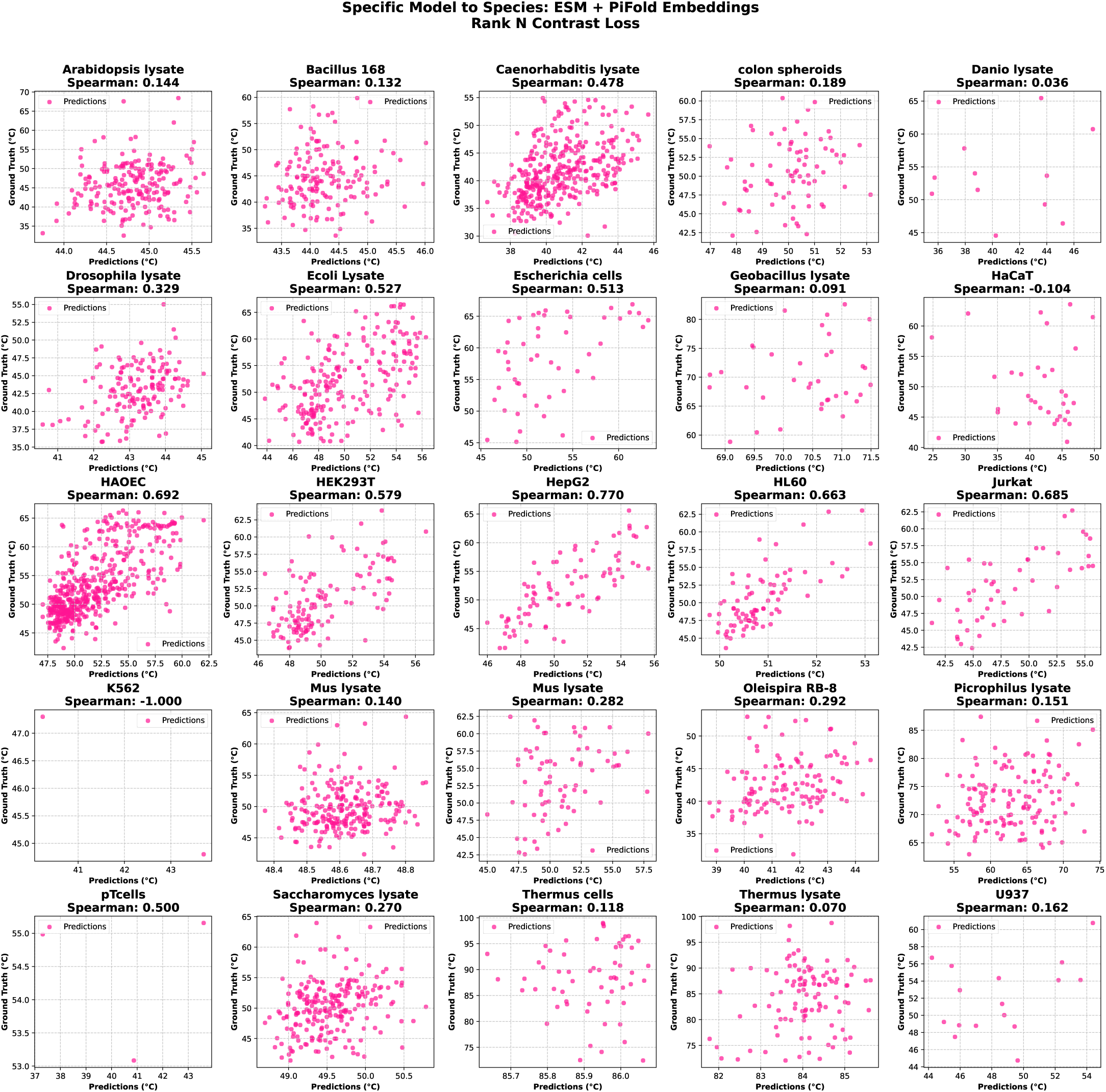
Scatterplot of individual models applied to their target species using Rank N Contrast loss in combination with ESM and PiFold embeddings.

## Notes

### Competing Interest Statement

The authors have declared no competing interest.

### Summary of Updates

This updated version provides a refined analysis of protein thermostability prediction, focusing on melting temperature. It builds on previous work by comparing two modeling approaches: transfer learning and Low-Rank Adaptation (LoRA) fine-tuning. These approaches were assessed in both cross-species (global) and species-specific settings, allowing for a more thorough evaluation of their ability to generalize across organisms and specialize within individual species. To strengthen the reliability of the results, statistical significance tests were included to support the observed differences between modeling strategies. Additional experiments were also conducted to confirm and expand upon the initial findings. Together, these improvements offer a more robust and detailed understanding of the trade-offs involved in different approaches to modeling protein thermostability.

